# Virus-derived peptide inhibitors of the herpes simplex virus type 1 nuclear egress complex

**DOI:** 10.1101/2020.06.24.168898

**Authors:** Elizabeth B. Draganova, Ekaterina E. Heldwein

## Abstract

Herpesviruses infect a majority of the human population, establishing lifelong latent infections for which there is no cure. Periodic viral reactivation spreads infection to new hosts while causing various disease states particularly detrimental in the immunocompromised. Efficient viral replication, and ultimately the spread of infection, is dependent on the nuclear egress complex (NEC), a conserved viral heterodimer that helps translocate mature viral capsids from the nucleus to the cytoplasm where they mature into infectious virions. Here, we have identified peptides capable of inhibiting the membrane-budding activity of the NEC *in vitro*. Biophysical characterization of these peptides using circular dichroism suggests that secondary structure, rather than amino acid sequence, influences the extent of inhibition. Current therapeutics that target viral DNA replication machinery are rendered ineffective by drug resistance due to viral mutations. Our results establish a basis for the development of an alternative class of inhibitors against nuclear egress, an essential step in herpesvirus replication, potentially expanding the current repertoire of available therapeutics.

---

The design of novel antiviral therapeutics is a rapidly expanding field because viruses remain one of the most common causes of human disease. One important group of viral human pathogens are herpesviruses that infects a majority of the population by establishing lifelong dormant infections for which there is no cure (reviewed in ^1–3^). Periodically, the virus reactivates, resulting in diseases such as keratitis, blindness, and cancers, which pose a significant threat to immunocompromised individuals. Current antiviral treatments of choice are nucleoside analogs such as acyclovir that target DNA replication machinery; yet, viral resistance to these drugs renders them ineffective, highlighting the need for additional therapeutics^4–6^.

The spread of infection to uninfected tissues and hosts relies on the ability of the virus to effectively replicate^2,7^. During replication, viral capsids are assembled within the nucleus and must translocate into the cytoplasm to form infectious virions but are too large to traverse nuclear pores, and instead use an unusual route termed nuclear egress. The first step in this process is the budding of a capsid at the inner nuclear membrane (INM), which is mediated by the conserved nuclear egress complex (NEC), a heterodimer of viral proteins UL31 and UL34^2,7,8^. The NEC is anchored to the INM by the single C-terminal transmembrane helix of UL34^9^. UL31 colocalizes with UL34^10,11^ so that the complex extends outward into the nucleoplasm where UL31 binds capsids ready for egress^12,13^.

Our group has demonstrated that purified recombinant NEC expressed in *E. coli* vesiculates synthetic membranes (giant unilamellar vesicles; GUVs) in the absence of other proteins or ATP^14^. Using cryo-electron microscopy and tomography (cryo-EM/ET), we showed that during budding, NEC oligomerizes into hexagonal membrane-bound coats^14^. We also found that HSV-1 NEC formed a hexagonal lattice of similar dimensions in crystals, which reinforced the inherent ability of the complex to form hexagonal arrays and provided an atomic-level view of protein interactions within the coat^15^. Mutations intended to disrupt hexagonal lattice formation impaired NEC-mediated budding *in vitro* and reduced nuclear egress and viral replication *in vivo*^16,17^, which confirmed that NEC oligomerization is essential for the NEC function. Furthermore, removal of NEC regions required for binding membranes *in vitro*^14^ also reduced viral replication *in vivo^18,19^*, suggesting that appropriate membrane interactions are also required for budding. Therefore, NEC-mediated budding is as an attractive therapeutic target.

Recently, we showed that HSV-1 NEC-mediated budding could be inhibited *in vitro* by another HSV-1 protein, capsid protein UL25^20^. UL25 is located at the capsid vertices and is thought to anchor the NEC to the capsid during nuclear egress^13,21^. We found that an N-terminally truncated UL25 construct UL25Δ44 Q72A (Fig. 1) inhibited NEC budding at a 10:1 molar ratio of UL25:NEC^20^. Using cryo-ET, we showed that the observed inhibitory effect on NEC budding correlated with a network of interconnected UL25 stars formed on top of the membrane-bound NEC layer. We hypothesized that this UL25 network blocked budding by preventing the conformational changes within the NEC necessary to generate membrane deformation and budding. UL25-mediated budding inhibition required the N-terminal α-helix in UL25, residues 45-94 (Fig. 1a), as a UL25Δ73 construct in which the N-terminal ~1/2 of this helix was truncated did not inhibit budding. We then tested whether peptides derived from the N-terminal α-helix of UL25 could inhibit NEC-mediated budding. Here, we show that UL25-derived peptides inhibit NEC budding and identify structural characteristics essential for inhibition.

**Figure 1.**
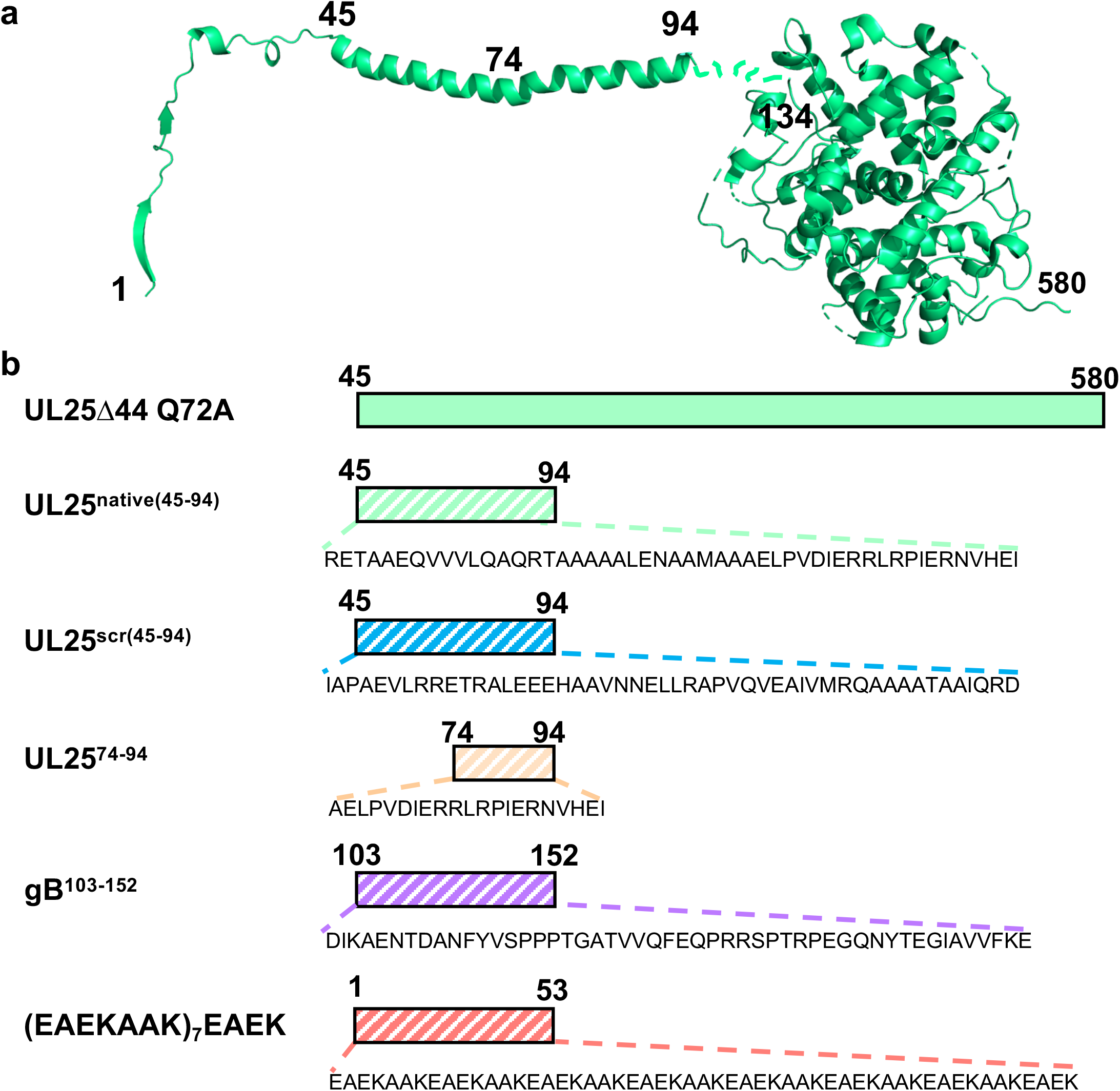
Design of UL25-derived peptides. **a)** The structure of the full-length, capsid-bound HSV-1 UL25 [residues 1-94 determined by cryo-EM (PDB ID: 6CGR)]; residues 134-580 determined by x-ray crystallography (PDB ID: 2F5U)]. The image of the UL25 structure was generated in PyMOL29. **b)** Schematic representation of peptides and protein constructs used in inhibition studies. Peptides are indicated by stripes. Amino-acid sequence of each peptide is shown. UL25Δ44 Q72A: N-terminally truncated UL25 mutant, used to yield pure protein; UL25^native(45-94)^: the native N-terminal helix of UL25; UL25^scr(45-94)^: the amino acid sequence of UL25^native(45-94)^ scrambled; UL25^74–94^: the C-terminal half of the UL25^native(45-94)^ peptide; gB^103–152^: a region within the HSV-1 glycoprotein B (gB); (EAEKAAK)_7_EAEK: a well-characterized α-helical peptide made of EAEKAAK repeats.

## Results

### UL25-derived peptides inhibit NEC budding, in a dose-dependent, length-dependent, but sequence-independent manner

We first tested whether the peptide corresponding to the entire N-terminal helix of UL25, UL25^native(45-94)^ (Fig. 1b), inhibited NEC-mediated budding in an established *in-vitro* GUV budding assay^14^ (Fig. 2a, b). NEC and UL25^native(45-94)^ were added to GUVs in 1:0.1, 1:0.3, 1:0.6, 1:1 and 1:10 molar ratios of NEC:peptide, and budding efficiency was quantified. We found that the UL25^native(45-94)^ peptide inhibited NEC budding in a dose-dependent manner and that the maximal inhibitory effect, a ~75% reduction in budding, was achieved at a 1:1 molar ratio of NEC:peptide (Fig. 2c). Surprisingly, a scrambled UL25^scr(45-94)^ peptide (Fig. 1b) inhibited budding with a similar efficiency (Fig. 2c), which suggested that the native UL25 peptide sequence was unimportant for the inhibitory activity. By contrast, a shorter UL25^74–94^ peptide (Fig. 1b) had a weak inhibitory effect on budding at a 1:1 or 1:10 molar ratios, ~25%, which was not statistically significant (Fig. 2c). Therefore, the inhibitory activity of the UL25-derived peptides is length-dependent.

**Figure 2.**
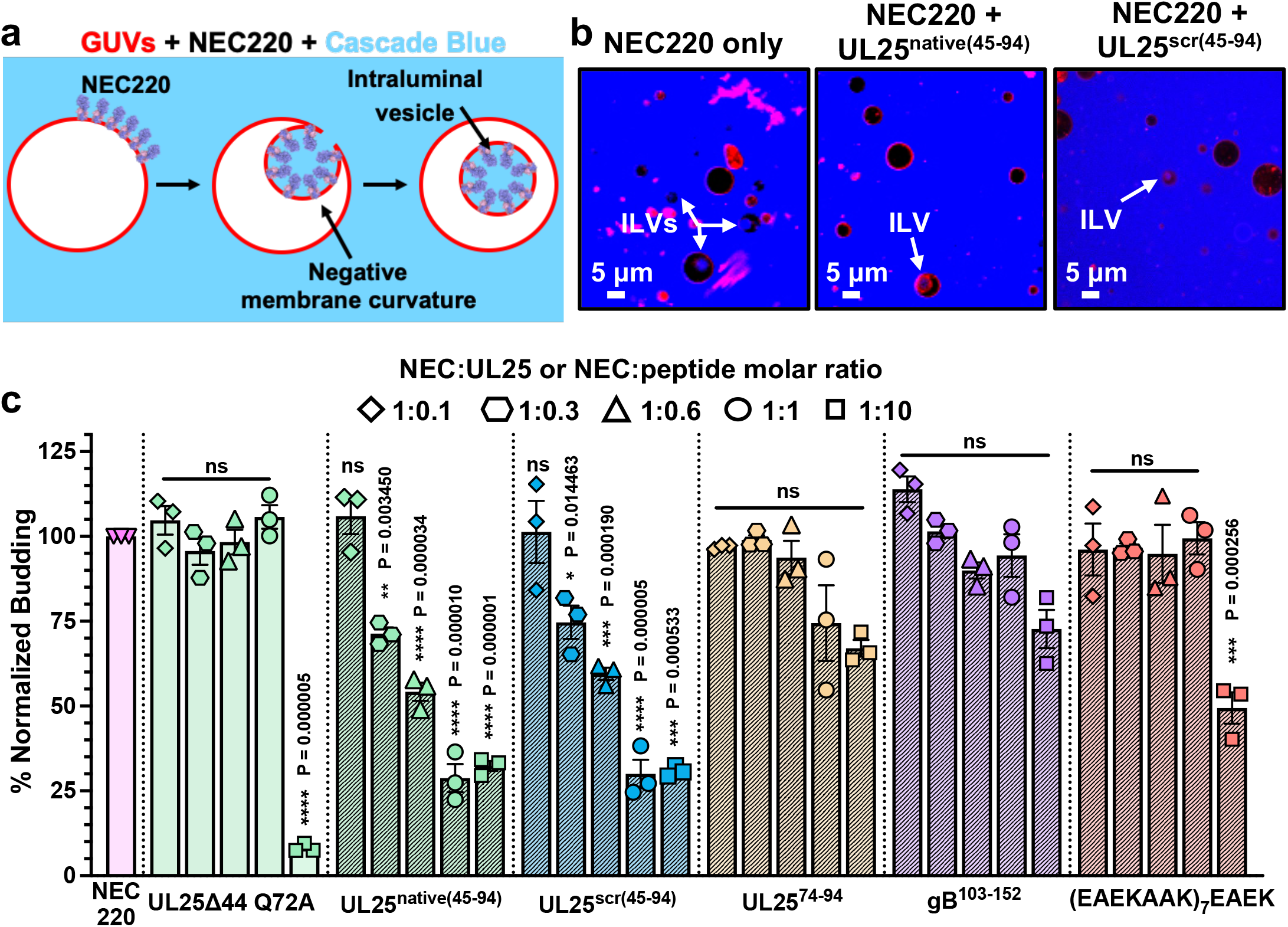
UL25^native(45-94)^ and UL25^scr(45-94)^ inhibit HSV-1 NEC-mediated budding in a dose-dependent manner. **a)** A diagram of the NEC-mediated *in-vitro* budding assay generated in Microsoft PowerPoint. The NEC [PDB ID: 4ZXS] binds red fluorescent giant unilamellar vesicles (GUVs) in the presence of the membrane impermeable Cascade Blue dye, inducing negative membrane curvature, and generates intraluminal vesicles (ILVs) filled with Cascade Blue. The number of ILV-containing GUVs serves as a measure of budding efficiency. **b)** Confocal microscopy images show ILVs formed inside the GUVs as the result of the NEC-mediated budding. Fewer ILVs are formed in the presence of UL25^native(45-94)^ or UL25^scr(45-94)^ due to inhibition of budding by the peptides. All images show the red (ATTO-DOPE) and blue (Cascade Blue) channels. Scale bars = 5 μm. **c)** Quantification of the NEC-mediated budding in the absence or the presence of peptides or UL25Δ44 Q72A. Budding was tested at 1:0.1, 1:0.3, 1:0.6, 1:1 or 1:10 molar ratios of NEC:UL25 or NEC:peptide. Each experiment was done in three biological replicates, each consisting of three technical replicates. Symbols represent average budding efficiency of each biological replicate relative to NEC220 (100%). Bars show the mean of three biological replicates, with error bars representing the standard error of the mean. Significance compared to NEC220 was calculated using an unpaired *t*-test. *P-value < 0.05, **P-value < 0.01, ***P-value < 0.001, and ****P-value < 0.0001.

UL25Δ44 Q72A, which served as a control, was not inhibitory at a 1:1 molar ratio (Fig. 2c), in agreement with our previous findings^20^; yet at a 1:10 molar ratio of NEC:UL25, it reduced NEC budding by ~90% (Fig. 2c). The differences in inhibition between UL25 and UL25-derived peptides are likely due to distinct inhibitory mechanisms.

### The ability of the peptides to form α-helical structure correlates with inhibition of NEC-mediated budding

Within capsid-bound UL25, amino acids 48-94 form an α-helix (Fig. 1a). Yet, the circular dichroism (CD) spectra of both the UL25^native(45-94)^ and the UL25^scr(45-94)^ peptides lacked any obvious α-helical signature in solution (Fig. 3a, b) and were only ~10% α-helical according to the estimates using DichroWeb^23^ (Supplementary Table S1). To assess the propensity of the UL25 peptides to form α-helical structure under certain conditions, the CD experiment was repeated in the presence of 30% trifluoroethanol (TFE), which is known to stabilize secondary structure^24^. In the presence of TFE, spectra of both peptides had clear α-helical structure (Fig. 3a, b), ~37% and ~31%, respectively (Supplementary Table S1), suggesting that each peptide could, in principle, form an α-helix in the proper environment, for example, when bound to a protein partner such as the NEC. The θ222/θ208 ratios of both peptides either in the absence or the presence of TFE were < 0.9 indicating peptides were forming individual helices rather than coiled coils^25^.

**Figure 3.**
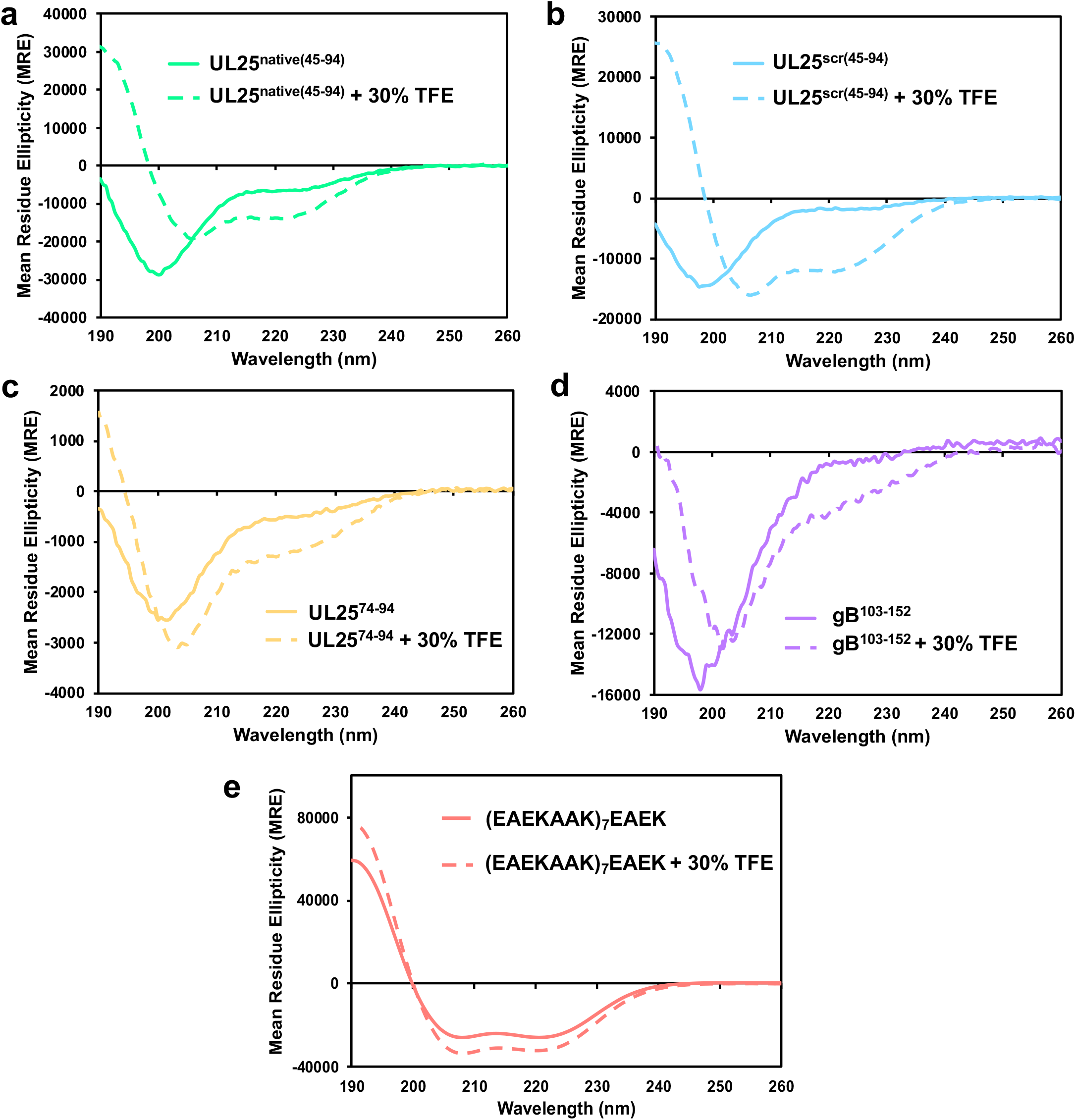
Secondary structure of the peptides used in this work. **a-e)** Far-UV CD spectra of each peptide in the presence of 30% TFE (dashed lines) or in its absence (solid lines). Both the UL25^native(45-94)^ and the UL25^scr(45-94)^ peptides form α-helices in the presence of 30% TFE. UL25^74-94^ and gB^103-152^ are mostly unstructured even in the presence of 30% TFE. The (EAEKAAK)_7_EAEK peptide is α-helical regardless of the presence of TFE.

The shorter UL25^scr(74-94)^ peptide adopted a random-coil conformation in solution even in the presence of 30% TFE (Fig. 3c, Supplementary Table 1). Therefore, it may be unable to inhibit NEC budding to a similar extent as the longer UL25^native(45-94)^ and the UL25^scr(45-94)^ peptides because it cannot form an α-helix.

### A pre-formed α-helical peptide moderately inhibits budding whereas a random-coil peptide does not

The inhibitory UL25-derived peptides can form α-helical structure whereas the non-inhibitory peptide does not, implying a correlation between the two properties. However, the inhibitory peptides are also substantially longer than the non-inhibitory one. Therefore, to explore the apparent correlation between the propensity to form an α-helix and the ability to inhibit NEC-mediated budding while ruling out the peptide length as a variable, we tested two heterologous peptides: a 50-amino-acid sequence from HSV-1 glycoprotein B (gB^103–152^), which lacks any regular secondary structure in the crystal structure^26^ and a 53-amino-acid peptide containing a series of well-characterized α-helical EAEKAAK repeats [(EAEKAAK)_7_EAEK]^27^ (Fig. 1b). The gB^103–152^ peptide was not inhibitory at a 1:1 molar ratio of NEC:peptide (Fig. 2c) and only had a weak inhibitory effect on budding at a 1:10 molar ratio of NEC:peptide, ~25%, which was not statistically significant (Fig. 2c). CD analysis showed this peptide adopted a random-coil conformation in solution even in the presence of 30% TFE (Fig. 3d, Supplementary Table 1). These observations suggest that the ability of the peptides to form an α-helix is important for their inhibitory activity.

The (EAEKAAK)_7_EAEK peptide was 60% α-helical in solution even in the absence of TFE (Fig. 3e, Supplementary Table 1), in contrast to both the UL25^native(45-94)^ and the UL25^scr(45-94)^ peptides that became mostly α-helical only in the presence of TFE (Fig. 3a, b). Yet, it was not inhibitory at a 1:1 molar ratio of NEC:peptide and moderately inhibited NEC-mediated budding at a 1:10 molar ratio, by ~50% (Fig. 2c). The differences in the inhibitory activities of the three peptides that can form α-helices could be due to different amino acid composition, especially, the charged residue content. The (EAEKAAK)_7_EAEK peptide contains 60% charged residues (glutamates and lysines) whereas the UL25-derived inhibitory peptides contain only 30% charged residues (aspartates, glutamates and arginines). The moderate inhibitory activity of the (EAEKAAK)_7_EAEK peptide at high concentrations could be due to its non-specific interactions with the NEC due to high charge. Alternatively, optimal inhibition may require a peptide that forms an α-helix in the presence of the NEC, perhaps, in an induced-fit manner, instead of a stable, pre-formed α-helix.

### Inhibitory peptides do not block NEC/membrane interactions

Peptides could inhibit NEC-mediated budding by preventing the NEC from binding to membranes. We first tested whether the peptides themselves could bind multilamellar vesicles (MLVs) of the same lipid composition as the GUVs used in the budding assay and found that none did (Fig. 4). This suggested that the peptides do not compete with the NEC for binding to membranes. We then tested whether the inhibitory peptides could block the NEC from binding to membranes.

**Figure 4.**
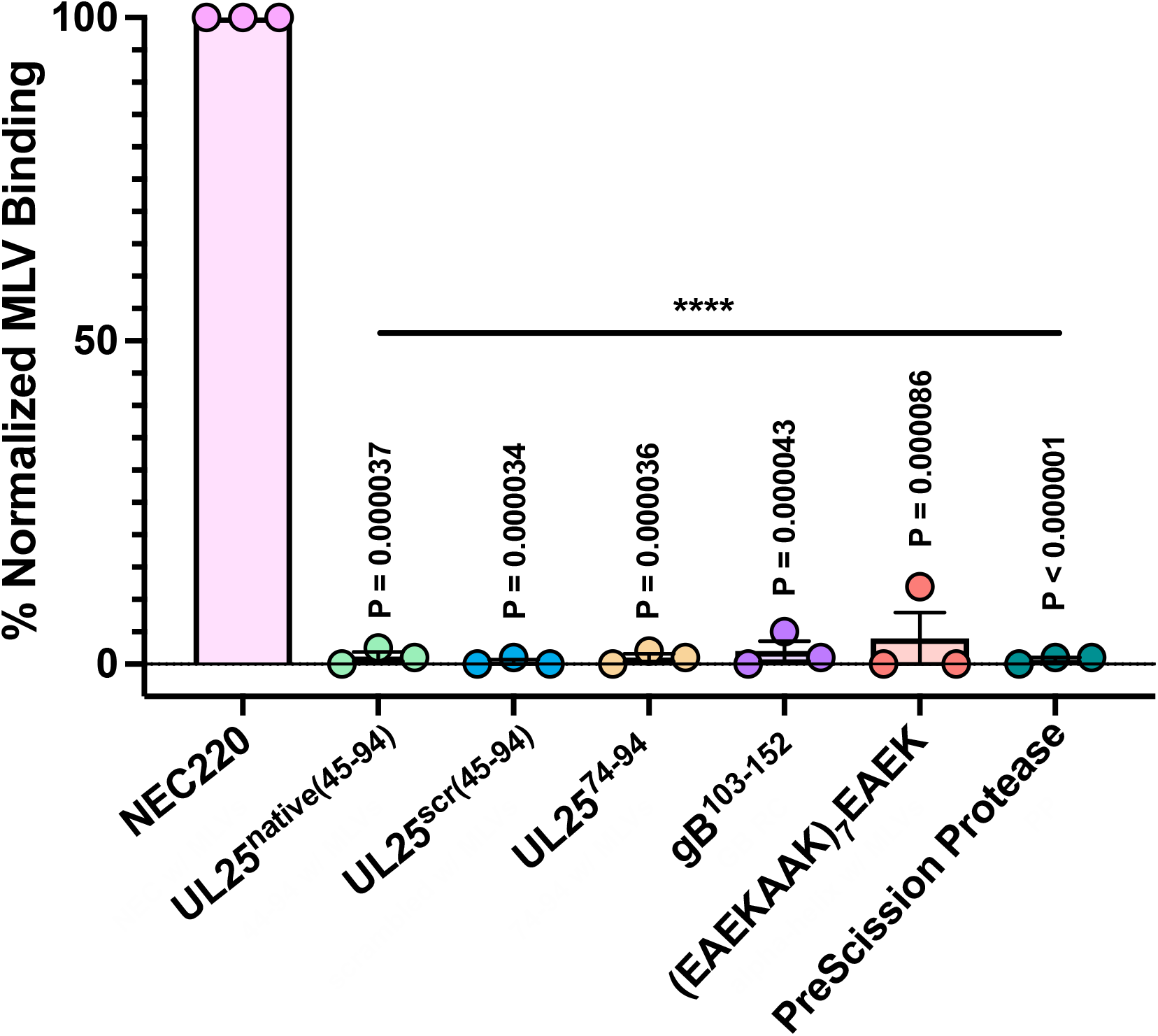
Peptides do not bind synthetic liposomes. NEC but not peptides bind multilamellar vesicles (MLVs) of the same composition as the GUVs used in the in-vitro budding assay. PreScission protease, which does not bind membranes, was used as a negative control. Each experiment was done in three biological replicates, each consisting of two technical replicates. Symbols represent average binding efficiency of each biological replicate relative to NEC220 (100%). Bars show the mean of three biological replicates, with error bars representing the standard error of the mean. Significance compared to NEC220 was calculated using an unpaired *t*-test. ****P-value < 0.0001.

To assess if the NEC could bind membranes in presence of inhibitory peptides, we performed a co-sedimentation assay with NEC, MLVs, and either UL25^native(45-94)^ or UL25^scr(45-94)^. UL25Δ44 Q72A, which inhibits budding but does not prevent the NEC from binding membranes^20^, was used as a control. Experiments were performed at a 1:10 molar ratio of NEC:peptide or NEC:UL25 molar ratio because the UL25Δ44 Q72A control inhibits at a 1:10 molar ratio of NEC:UL25. We found that neither peptide prevented the NEC from binding membranes as judged by the same amount of MLV-bound NEC (~95%) in the absence or the presence of peptides (Fig. 5). As expected^20^, UL25Δ44 Q72A also did not block NEC membrane binding (Fig. 5). Thus, the inhibitory peptides do not block NEC/membrane interactions and, instead, inhibit NEC budding by some other mechanism.

**Figure 5.**
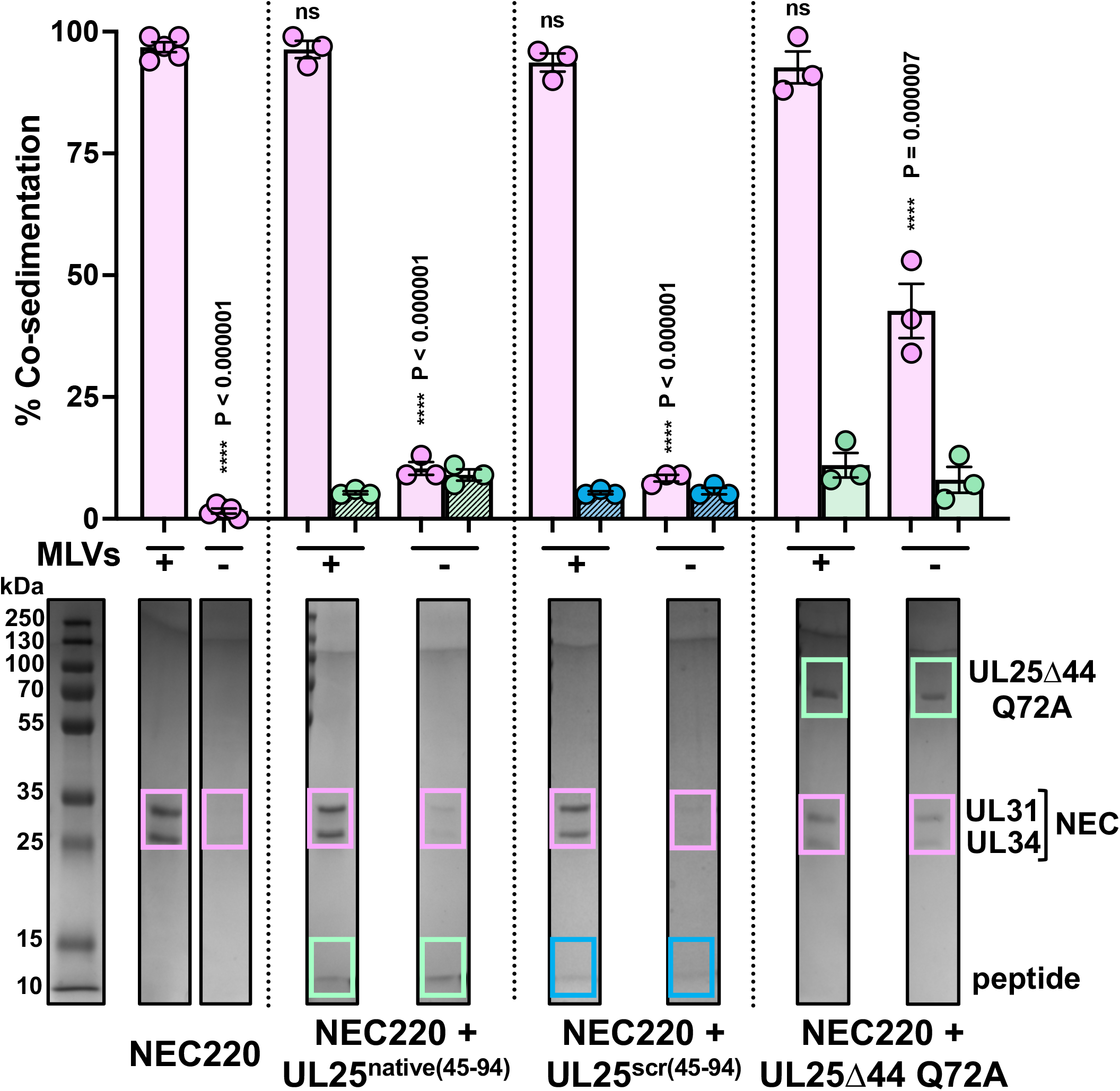
Peptides do not block membrane binding by the NEC. Co-sedimentation of NEC220 alone and in the presence of either UL25^native(45-94)^, UL25^scr(45-94)^, or UL25Δ44 Q72A was assessed by a co-sedimentation assay in the presence and absence of MLVs. The amount of co-sedimentation was quantified with Coomassie staining. Bands from representative gels (full gel images are shown in Supplementary Figure 1) used for quantification are boxed and colored as follows: pink – NEC (UL31: 34 kDa and UL34: 25 kDa); green – either UL25^native(45-94)^ (5 kDa) or UL25Δ44 Q72A (57 kDa); blue – UL25^scr(45-94^) (5 kDa). Each experiment was done in three biological replicates, each consisting of two technical replicates. Symbols represent average binding efficiency of each biological replicate. Bars show the mean of three biological replicates, with error bars representing the standard error of the mean. Significance compared to NEC220 in the presence of MLVs was calculated using an unpaired *t*-test. ****P-value < 0.0001.

## Discussion

Here, we have shown that two peptides derived from the N-terminal α-helix of UL25 efficiently inhibited NEC-mediated budding *in vitro* in a dose-dependent manner. The inhibitory ability of the peptides depended on their length and the ability to form an α-helix but not on the exact amino acid sequence. Peptide length was more important for inhibition than the sequence because 50-amino-acid UL25 peptides, be they native or scrambled (UL25^native(45-94)^ and UL25^scr(45-94)^), efficiently inhibited budding whereas the 21-amino-acid native peptide UL25^74–94^ did not. Future studies will identify the minimal UL25 peptide length required for inhibition. The inhibitory ability of UL25 peptides correlated not only with their length but also with the propensity to adopt α-helical structure. Although all three UL25-derived peptides formed random coils in solution, the two 50-amino-acid peptides formed α-helices in the presence of TFE, known to stabilize secondary structure, but the 21-amino-acid native peptide did not.

To uncouple the propensity to form α-helices from the peptide length, we tested a heterologous 50-amino-acid peptide from HSV-1 gB (gB^103–152^) that lacks any regular secondary structure in the crystal structure^26^. We confirmed that this peptide formed a random coil and showed that had a weak inhibitory activity only at high molar ratios of NEC:peptide.

We also tested a heterologous 53-amino-acid peptide containing a series of well-characterized α-helical EAEKAAK repeats [(EAEKAAK)_7_EAEK]^27^. We found that, unlike the inhibitory UL25-derived peptides, this peptide was α-helical in solution even in the absence of TFE and that it had a modest inhibitory effect only at the highest molar ratio of NEC:peptide. This heterologous α-helical peptide is highly charged and forms an amphipathic α-helix with a charged face rich in glutamates and lysines and a hydrophobic face rich in alanines (Supplementary Fig. S2). In contrast, inhibitory UL25-derived peptides are less charged and form α-helices with a less pronounced amphipathic character (Supplementary Fig. S2). The moderate inhibitory activity of the (EAEKAAK)_7_EAEK peptide at high concentrations could be due to its non-specific interactions with the NEC. Alternatively, it is possible that a stable, preformed α-helix may not be able to form appropriate inhibitory contacts with the NEC and that for efficient inhibition, an α-helix of a specific length and amino-acid content should form in the presence of the NEC, in an induced-fit manner. Therefore, future design of NEC peptide inhibitors should take both of these characteristics into consideration.

The mechanism by which the inhibitory peptides inhibit budding is yet unclear. Our data suggest that the inhibitory peptides do not prevent the NEC from binding the membranes. Therefore, we hypothesize that the peptides may, instead, inhibit the membrane-budding activity of the NEC by some other mechanism, for example, by blocking NEC/NEC interactions required for budding or prevent conformational changes required by the NEC to undergo budding (Fig. 6).

**Figure 6.**
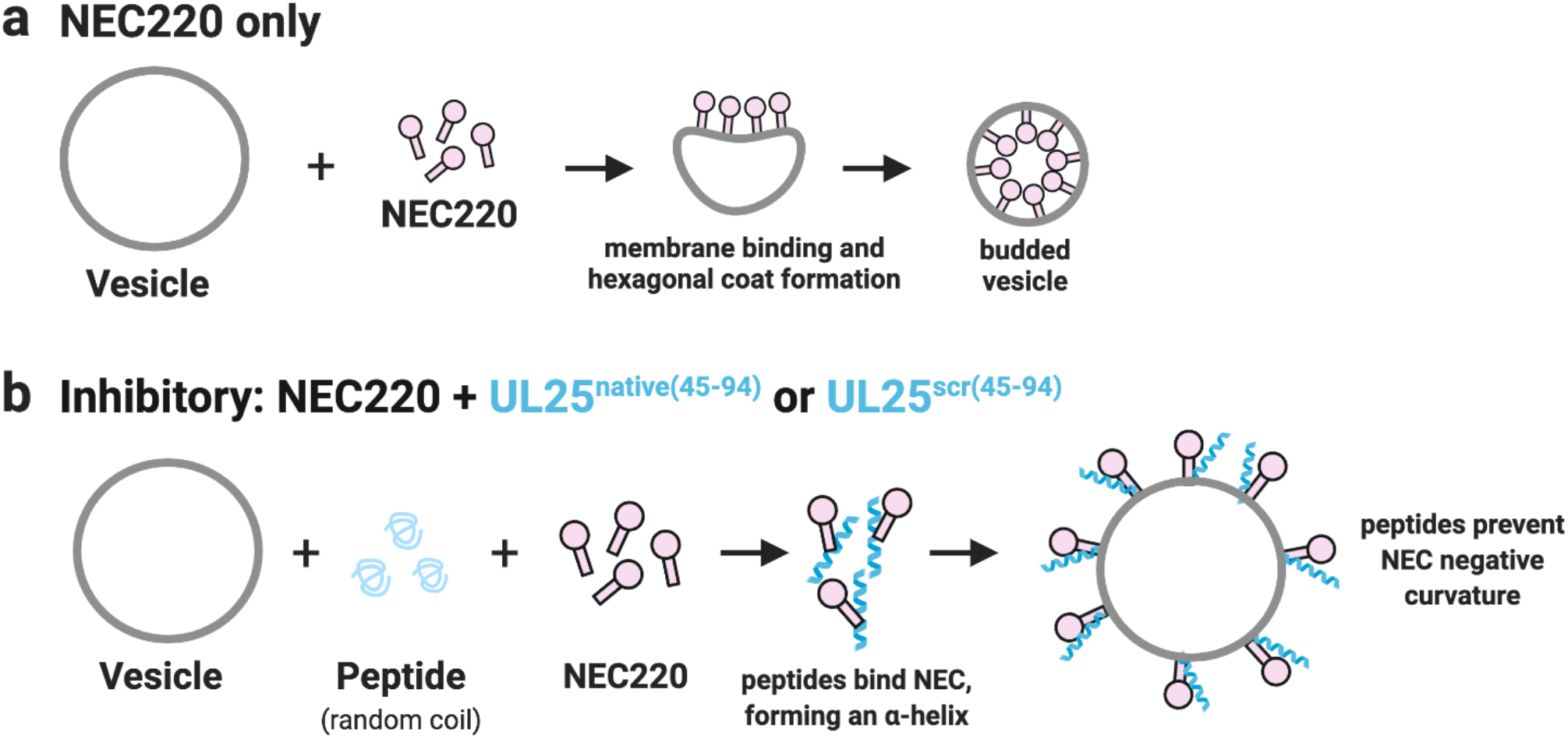
Proposed model of peptide inhibition of the NEC-mediated budding. **a)** NEC (pink) alone binds membranes, induces negative membrane curvature, and generates a budded vesicle containing an internal hexagonal NEC coat. **b)** Both UL25^native(45-94)^ and UL25^scr(45-94)^ (bluegreen) inhibit budding by forming α-helices upon binding to the NEC, which likely prevents conformational changes necessary for the formation of the negative membrane curvature. This figure was generated using Biorender (https://biorender.com).

Our previous study suggested that UL25 inhibited NEC-mediated budding by oligomerizing into a network of interconnected stars on top of the membrane-bound NEC layer as observed by cryo-ET^20^. We hypothesized that the N-terminal helices link the neighboring UL25 molecules into a network by forming antiparallel helical bundles that sterically blocked the NEC from undergoing conformational changes necessary for membrane budding. Both the N-terminal helix and the C-terminal core of UL25 (Fig. 1a) were important for inhibition. However, the inhibitory UL25 peptides in our current study were composed of only the N-terminal helix.

Therefore, they likely bind the NEC differently from UL25 and use a different mechanism to inhibit budding. Being much smaller than UL25, the inhibitory peptides can interact with the NEC at sites otherwise inaccessible to the UL25 protein. Alternatively, NEC/peptide interactions may block NEC/NEC interactions necessary for the assembly of the hexagonal NEC coat thereby blocking the induction of negative curvature and, ultimately, budding (Fig. 6). Additional experiments, such as fluorescent labeling of the peptides, are necessary to show peptide binding to membrane-bound NEC.

Our results suggest that inhibitory peptides should be of a certain length, moderately charged, and unstructured in solution, yet capable of forming α-helical structure, presumably, upon binding to the NEC. Further structural analyses of the inhibitory peptides in the presence of the NEC are necessary to narrow down the optimal characteristics of the inhibitory peptides, including their delivery to their intracellular target. The length of the inhibitory peptides identified in this study, 50 amino acids, presents challenges for intracellular delivery. Most therapeutic peptides are ~20-30 amino acids in length, which increases the likelihood of crossing the plasma membrane, yet many require additional alterations of their physical properties to do so^28^. Furthermore, peptides are susceptible to proteolytic degradation and can exhibit off-target effects in cells, both of which the likelihood increasing the longer the peptide, reducing the chances of binding the intended target. Therefore, determining the minimal inhibitory length of these peptides will be essential for transitioning from *in-vitro* to *in-vivo* studies. Lastly, the NEC is located within the nucleus, requiring inhibitory peptides to not only reach the cytoplasm but also enter the nucleus. A recent peptide cellular localization assay showed that peptides smaller than 40 kDa can diffuse through the nuclear pore^29^, suggesting NEC inhibitory peptides could do so as well.

Efficient NEC function is required for successful replication of herpesviruses. To our knowledge, this is the first study reporting a peptide-based approach targeting NEC budding. Our work provides a starting point for the rational design of peptide-based inhibitors of the NEC and nuclear egress. Both the NEC and UL25 are conserved across all herpesviruses raising an intriguing possibility that UL25-derived peptides could be used to inhibit NEC homologs from other herpesviruses.

## Materials and Methods

### Peptides

All peptides used in this study were purchased from Peptide 2.0 with the exception of the α-helical (EAEKAAK)_7_EAEK peptide, which was cloned, expressed and purified as described below.

### Cloning

All primers used in cloning are listed in Supplementary Table S2. The DNA sequence encoding the α-helical [(EAEKAAK)_7_EAEK] peptide was subcloned into the prokaryotic expression vector pGEX-6P-1 that encodes a N-terminal GST-tag followed by a PreScission Protease cleavage site in frame with the BamHI restriction site within the multiple cloning site. The (EAEKAAK)_7_EAEK peptide DNA sequence was obtained as a gBlock gene fragment (IDT) and subjected to restriction digest cloning with BamHI and XhoI into the pGEX-6P-1 vector (the gene fragment sequence is listed in Supplementary Table S2) to create the pED39 [(EAEKAAK)_7_EAEK peptide] plasmid.

### Expression and purification of NEC220

Plasmids encoding HSV-1 UL31 1-306 (pKH90) and UL34 1-220 (pJB02) were co-transformed into *Escherichia coli* BL21(DE3) LoBSTr cells (Kerafast) to generate NEC220 (NEC construct absent of the UL34 C-terminal transmembrane helix)^14^. NEC220 was expressed using autoinduction at 37 °C in Terrific Broth (TB) supplemented with 100 μg/mL kanamycin, 100 μg/mL ampicillin, 0.2% lactose, and 2 mM MgSO4 for 4 h. The temperature was then reduced to 25 °C for 16 h. Cells were harvested at 5,000 × g for 30 min. NEC220 was purified as previously described^14^ with slight modifications. NEC220 was passed over 2 × 1 mL HiTrap Talon columns (GE Healthcare), rather than ion exchange, to remove excess cleaved His6-SUMO before injection onto size-exclusion chromatography.

### Expression and purification of UL25Δ44 Q72A

The plasmid encoding HSV-1 UL25Δ44 Q72A (pED03) was expressed and cells harvested as described above for NEC220 except that only 100 μg/mL kanamycin was used. UL25Δ44 Q72A was purified as previously described^20^.

### Expression and purification of (EAEKAAK)_7_EAEK

The plasmid encoding the peptide sequence was expressed and cells harvested as described above for the NEC220 with the exception that only 100 μg/mL ampicillin was used. Cells were resuspended in lysis buffer (50 mM Na HEPES pH 7.5, 500 mM NaCl, 1 mM TCEP, and 10% glycerol) plus Complete protease inhibitor (Roche) and lysed using a microfluidizer 110S (Microfluidics). The cell lysate was centrifuged at 13,000 × g for 35 min, and the supernatant passed over a Glutathione sepharose 4B (GE Healthcare) column, which was subsequently washed with lysis buffer. The GST-tag was cleaved on the glutathione sepharose column for 16 h using the PreScission Protease described above. The peptide was eluted off the column with lysis buffer. As a final purification step, the peptide was purified with size-exclusion chromatography using a Superdex 75 column (GE Healthcare) equilibrated with gel filtration buffer (20 mM Na HEPES, pH 7.0, 100 mM NaCl, and 1 mM TCEP) for budding assays or with CD buffer (10 mM sodium phosphate, pH 7.4, and 100 mM NaF) for CD studies. Absorbance was monitored at 214 nm because the peptide lacks aromatic residues. The peptide was purified to homogeneity as assessed by both 16.5% Tris-Tricine SDS-PAGE and 12% SDS-PAGE with Coomassie staining. Fractions containing peptide were concentrated and sent off for amino acid analysis (University of Colorado Denver Anschutz Medical Campus) for accurate concentration determination.

### Co-sedimentation assays

Co-sedimentation of either peptides alone or NEC and peptides (1:10 molar ratio of NEC:peptide) to multilamellar vesicles (MLVs) was performed as previously described^14^. MLVs were prepared in a 3:1:1 molar ratio of 1-palmitoyl-2-oleoyl-glycero-3-phosphocholine (POPC):1-palmitoyl-2-oleoyl-sn-glycero-3-phospho-L-serine (POPS):1-palmitoyl-2-oleoyl-sn-glycero-3-phosphate (POPA) (Avanti Polar Lipids). Background signal in the absence of liposomes is due to protein aggregation during centrifugation. Each experiment was done in three biological replicates with two technical replicates. For peptides alone, the reported values represent the average binding of each biological replicate of peptide to MLVs relative to the NEC (100%). For experiments measuring NEC/MLV binding in the presence of peptides, reported values represent the percentage (0-100%) of co-sedimentation of either NEC220 or peptides in the absence or presence of MLVs. The standard error of the mean is reported for each measurement.

### GUV Budding Assays

Giant unilamellar vesicles (GUVs) were prepared as previously described^14^. For NEC220 only budding quantification, a total of 10 μL of GUVs with a 3:1:1 molar ratio of POPC:POPS:POPA containing ATTO-594 DOPE (ATTO-TEC GmbH) at a concentration of 0.2 μg/μL was mixed with 1 μM NEC220 (final concentration), and 0.2 mg/mL (final concentration) Cascade Blue Hydrazide (ThermoFisher Scientific). For the NEC and peptide experiments, 10 μL of GUVs and 0.1, 0.3, 0.6, 1, or 10 μM (final concentration) of either the UL25^native(45-94)^, UL25^scr(45-94)^, UL25^74–94^, gB^103–152^, or (EAEKAAK)_7_EAEK peptides were incubated with 1 μM of NEC220 (final concentration) along with Cascade blue. The total volume of each sample during imaging for all experiments was brought to 100 μL with the gel filtration buffer and the reaction was incubated for 5 min at 20 °C. Samples were imaged in a 96-well chambered cover-glass. Images were acquired using a Nikon A1R Confocal Microscope with a 60x oil immersion lens at the Tufts Imaging Facility in the Center for Neuroscience Research at Tufts University School of Medicine. Quantification was performed by counting vesicles in 15 different frames of the sample (~300 vesicles total). Each experiment was done in at least three biological replicates, each with three technical replicates. Prior to analysis, the background was subtracted from the raw values. All raw values are given in Supplementary Table S3. The reported values represent the average budding activity relative to NEC220 (100%). The standard error of the mean is reported for each measurement. Significance compared to NEC220 was calculated using an unpaired one-tailed *t*-test against NEC220.

### Circular dichroism (CD) studies

Far-UV CD spectra of peptides (0.1 mg/mL) were recorded in 10 mM Na phosphate, pH 7.4, and 100 mM NaF buffer using a Jasco 815 CD Spectropolarimeter at the Center for Macromolecular Interactions at Harvard Medical School. Peptide spectra were also measured in the presence of 30% trifluoroethanol (TFE). Data were collected at ambient temperature with a scan speed of 50 nm/min and 5 accumulations of each sample was averaged. The raw data was blank subtracted and converted to mean residue ellipticity (MRE). Helical content was estimated using DichroWeb^23^, and average values from three reference datasets generated by the same analysis program are provided in Supplementary Table S1.

## Acknowledgments

We would like to thank Shaun Bevers (University of Colorado Denver Biophysics Core) for performing amino acid analysis on the peptides. We also thank Peter Cherepanov (Francis Crick Institute) for the gift of the GST-PreScission protease expression plasmid and Thomas Schwartz (Massachusetts Institute of Technology) for the gift of LoBSTr cells. CD experiments were performed at the Center for Macromolecular Interactions in the Department of Biological Chemistry and Molecular Pharmacology at Harvard Medical School. This work was funded by the NIH grants R01GM111795 and R01AI147625 (E.E.H.), a Faculty Scholar grant 55108533 from Howard Hughes Medical Institute (E.E.H.), NIH postdoctoral fellowship F32GM126760 (E.B.D.), and NIH IRACDA postdoctoral fellowship K12GM133314 (E.B.D).

## Author Contributions

E.B.D. and E.E.H. designed the experiments; E.B.D. performed all experiments and analyses under the guidance of E.E.H; E.B.D. and E.E.H. wrote the manuscript.

## References

1. Johnson, D. C. & Baines, J. D. Herpesviruses remodel host membranes for virus egress. Nat. Rev. Microbiol. 9, 382–394, doi:10.1038/nrmicro2559 (2011).

2. Bigalke, J. M. & Heldwein, E. E. Nuclear exodus: Herpesviruses lead the way. Annu. Rev. Virol. 3, 387–409, doi:10.1146/annurev-virology-110615-042215 (2016).

3. Mettenleiter, T. C., Klupp, B. G. & Granzow, H. Herpesvirus assembly: an update. Virus Res. 143, 222–234, doi:10.1016/j.virusres.2009.03.018 (2009).

4. Szczubialka, K., Pyrc, K. & Nowakowska, M. In search for effective and definitive treatment of herpes simplex virus type 1 (HSV-1) infections. RSC Adv. 6, 1058–1075, doi:10.1039/c5ra22896d (2016).

5. Engel, J. P., Englund, J. A., Fletcher, C. V. & Hill, E. L. Treatment of resistant herpes simplex virus with continuous-infusion acyclovir. JAMA. 263, 1662–1664 (1990).

6. Antoine, T. E., Park, P. J. & Shukla, D. Glycoprotein targeted therapeutics: a new era of anti-herpes simplex virus-1 therapeutics. Rev. Med. Virol. 23, 194–208, doi:10.1002/rmv.1740 (2013).

7. Draganova, E. B., Thorsen, M. K. & Heldwein, E. E. Nuclear Egress. Curr. Issues Mol. Biol. 41, 125–170, doi:10.21775/cimb.041.125 (2020).

8. Reynolds, A. E., Wills, E. G., Roller, R. J., Ryckman, B. J. & Baines, J. D. Ultrastructural localization of the herpes simplex virus type 1 UL31, UL34, and US3 proteins suggests specific roles in primary envelopment and egress of nucleocapsids. J. Virol. 76, 8939–8952 (2002).

9. Shiba, C. et al. The UL34 gene product of herpes simplex virus type 2 is a tail-anchored type II membrane protein that is significant for virus envelopment. J. Gen. Virol. 81, 2397–2405, doi:10.1099/0022-1317-81-10-2397 (2000).

10. Reynolds, A. E. et al. U(L)31 and U(L)34 proteins of herpes simplex virus type 1 form a complex that accumulates at the nuclear rim and is required for envelopment of nucleocapsids. J. Virol. 75, 8803–8817 (2001).

11. Chang, Y. E. & Roizman, B. The product of the UL31 gene of herpes simplex virus 1 is a nuclear phosphoprotein which partitions with the nuclear matrix. J. Virol. 67, 6348–6356 (1993).

12. Trus, B. L. et al. Allosteric signaling and a nuclear exit strategy: binding of UL25/UL17 heterodimers to DNA-Filled HSV-1 capsids. Mol. Cell. 26, 479–489, doi:10.1016/j.molcel.2007.04.010 (2007).

13. Yang, K. & Baines, J. D. Selection of HSV capsids for envelopment involves interaction between capsid surface components pUL31, pUL17, and pUL25. Proc. Natl. Acad. Sci. 108, 14276–14281, doi:10.1073/pnas.1108564108 (2011).

14. Bigalke, J. M., Heuser, T., Nicastro, D. & Heldwein, E. E. Membrane deformation and scission by the HSV-1 nuclear egress complex. Nat. Commun. 5, 4131, doi:10.1038/ncomms5131 (2014).

15. Bigalke, J. M. & Heldwein, E. E. Structural basis of membrane budding by the nuclear egress complex of herpesviruses. EMBO J. 34, 2921–2936, doi:10.15252/embj.201592359 (2015).

16. Roller, R. J., Zhou, Y., Schnetzer, R., Ferguson, J. & DeSalvo, D. Herpes simplex virus type 1 U(L)34 gene product is required for viral envelopment. J. Virol. 74, 117–129 (2000).

17. Fuchs, W., Klupp, B. G., Granzow, H., Osterrieder, N. & Mettenleiter, T. C. The interacting UL31 and UL34 gene products of pseudorabies virus are involved in egress from the host-cell nucleus and represent components of primary enveloped but not mature virions. J. Virol. 76, 364–378 (2002).

18. Mou, F., Wills, E. & Baines, J. D. Phosphorylation of the U(L)31 protein of herpes simplex virus 1 by the U(S)3-encoded kinase regulates localization of the nuclear envelopment complex and egress of nucleocapsids. J. Virol. 83, 5181–5191, doi:10.1128/jvi.00090-09 (2009).

19. Passvogel, L., Trube, P., Schuster, F., Klupp, B. G. & Mettenleiter, T. C. Mapping of sequences in Pseudorabies virus pUL34 that are required for formation and function of the nuclear egress complex. J. Virol. 87, 4475–4485, doi:10.1128/jvi.00021-13 (2013).

20. Draganova, E. B., Zhang, J., Zhou, Z. H. & Heldwein, E. E. Structural basis for capsid recruitment and coat formation during HSV-1 nuclear egress. Elife. 9, doi:10.7554/eLife.56627 (2020).

21. Yang, K., Wills, E., Lim, H. Y., Zhou, Z. H. & Baines, J. D. Association of herpes simplex virus pUL31 with capsid vertices and components of the capsid vertex-specific complex. J. Virol. 88, 3815–3825, doi:10.1128/jvi.03175-13 (2014).

22. DeLano, W., L. The PyMOL Molecular Graphics System, Version 1.7.4 Schrödinger, LLC. http://www.pymol.org (2015).

23. Whitmore, L. & Wallace, B. A. Protein secondary structure analyses from circular dichroism spectroscopy: methods and reference databases. Biopolymers. 89, 392–400, doi:10.1002/bip.20853 (2008).

24. Luo, P. & Baldwin, R. L. Mechanism of helix induction by trifluoroethanol: a framework for extrapolating the helix-forming properties of peptides from trifluoroethanol/water mixtures back to water. Biochemistry. 36, 8413–8421, doi:10.1021/bi9707133 (1997).

25. Lau, S. Y., Taneja, A. K. & Hodges, R. S. Synthesis of a model protein of defined secondary and quaternary structure. Effect of chain length on the stabilization and formation of two-stranded alpha-helical coiled-coils. J. Biol. Chem. 259, 13253–13261 (1984).

26. Heldwein, E. E. et al. Crystal structure of glycoprotein B from herpes simplex virus 1. Science. 313, 217–220, doi:10.1126/science.1126548 (2006).

27. Zhou, N. E., Kay, C. M., Sykes, B. D. & Hodges, R. S. A single-stranded amphipathic alpha-helix in aqueous solution: design, structural characterization, and its application for determining alpha-helical propensities of amino acids. Biochemistry. 32, 6190–6197 (1993).

28. Derakhshankhah, H. & Jafari, S. Cell penetrating peptides: A concise review with emphasis on biomedical applications. Biomed. Pharmacother. 108, 1090–1096, doi:10.1016/j.biopha.2018.09.097 (2018).

29. Peraro, L. et al. Cell Penetration Profiling Using the Chloroalkane Penetration Assay. J. Am. Chem. Soc. 140, 11360–11369, doi:10.1021/jacs.8b06144 (2018).

