## Supplemental Figures S1-S2 and Supplemental Tables S1-S3 for "Virus-derived peptide inhibitors of the herpes simplex virus type 1 nuclear egress complex"

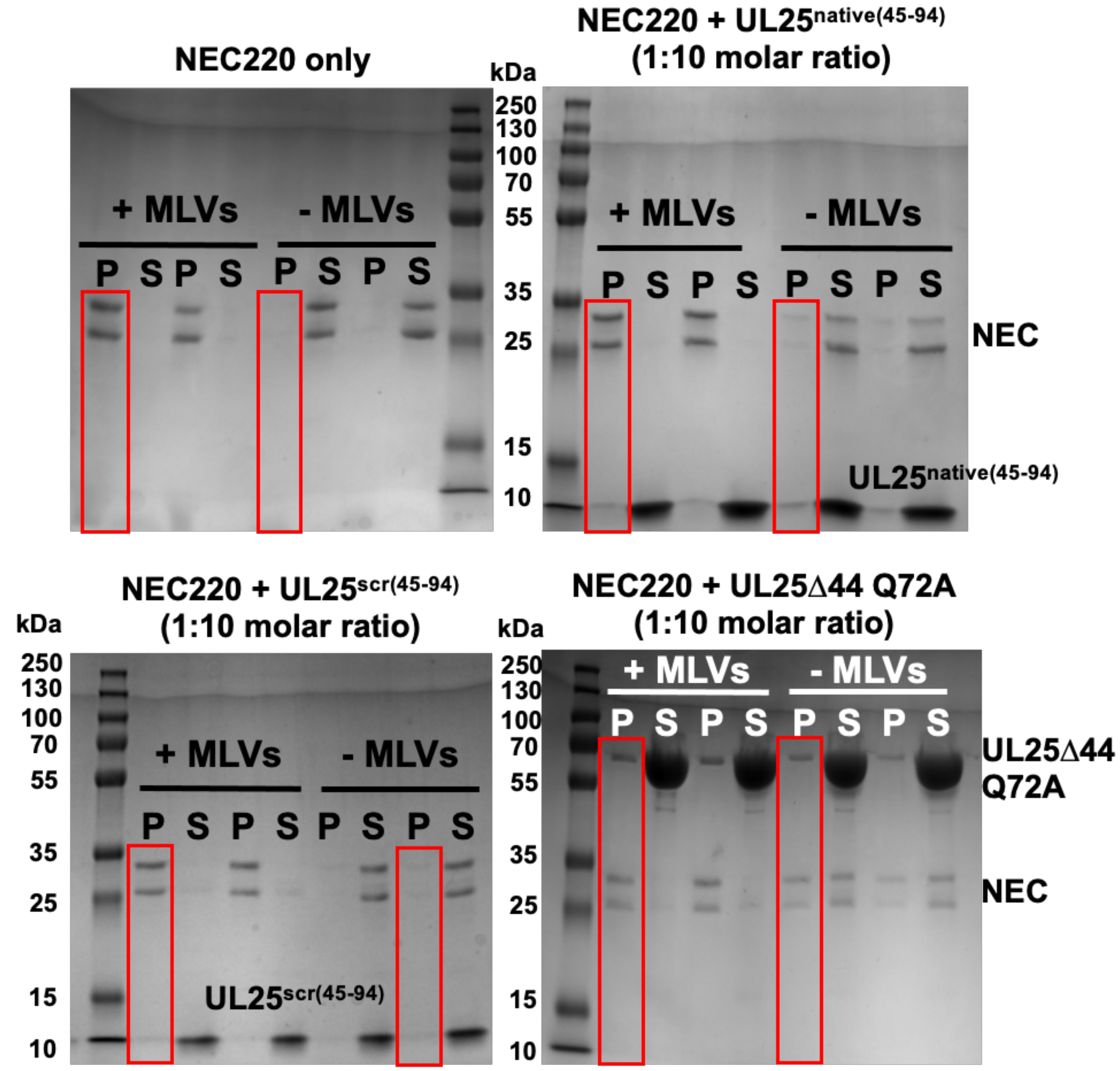


**Supplementary Fig. S1. Original SDS-PAGE images of the co-sedimentation assay samples shown in Figure 5.** Each gel represents one biological replicate used to generate the bar graph in Figure 5. All samples were run in parallel. Red boxes denote the gel lanes shown in the main figure. Gels were stained with Coomassie G-250. Gels were imaged together using a Syngene G:BOX. Both the pellet (P) and supernatant (S) samples, either in the presence (+MLVs) or absence (-MLVs) of MLVs are shown on the gel.

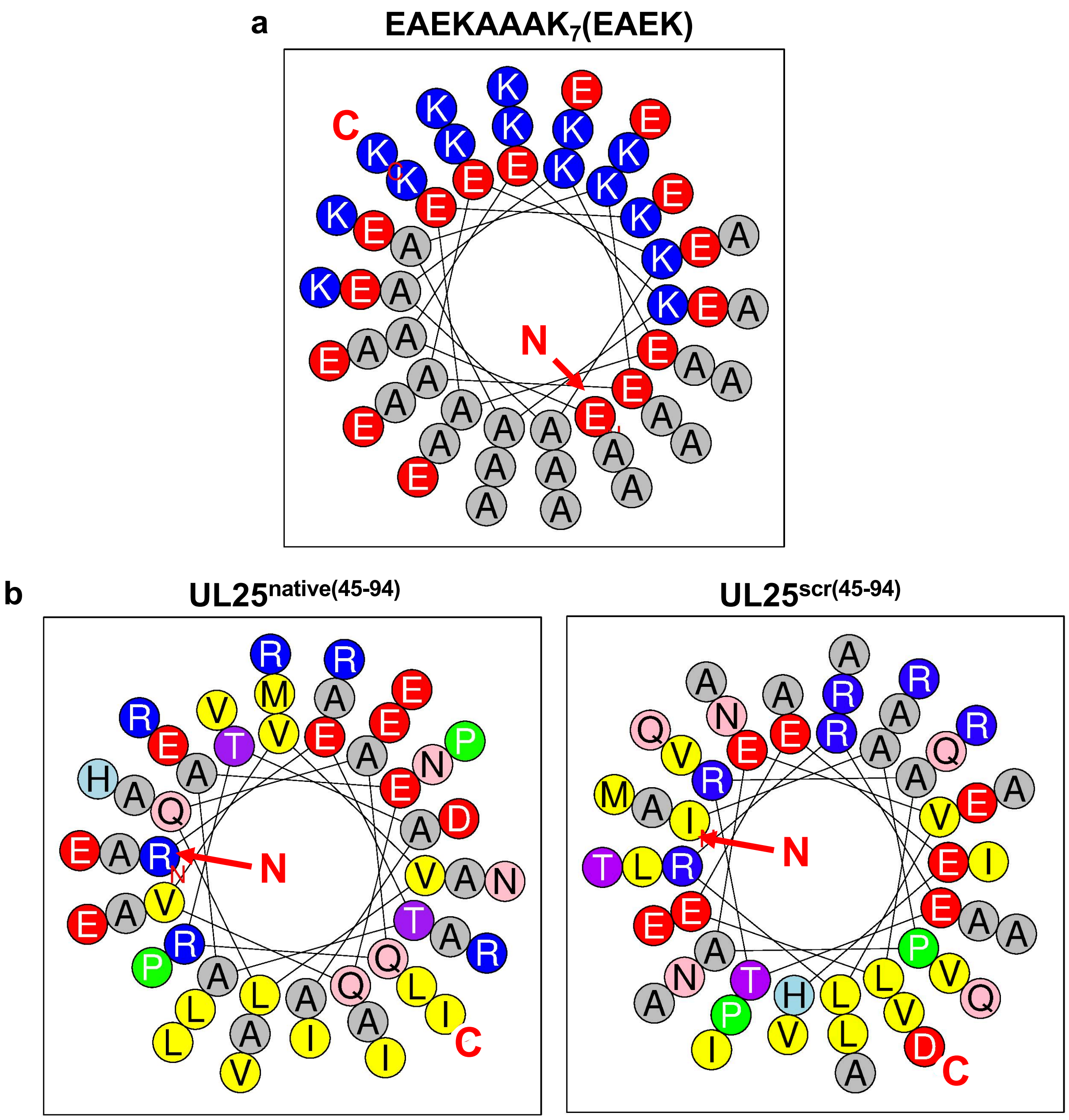


**Supplementary Fig. S2. Helical wheel projections of the (EAEKAAAK)$_7$EAEK, UL25$^{native(45-94)}$ and UL25$^{scr(45-94)}$ peptides. a)** The (EAEKAAAK)$_7$EAEK peptide is amphipathic as shown by the distribution of charged (blue and red) residues on one face of the helix versus the hydrophobic alanine residues (grey) on the other. **b)** The UL25$^{native(45-94)}$ and UL25$^{scr(45-94)}$ peptides are less amphipathic as indicated by the less even distribution of charged (blue and red) and polar residues (pink, light blue, and purple) versus hydrophobic (grey and yellow) residues. The N and C on each wheel indicate the N and the C termini. Wheel projections were made using Heliquest[1].

**Supplementary Table S1. Secondary structure content (%) estimation of peptides**. Spectral deconvolution using DichroWeb[2] was performed on samples in the absence or presence of 30% trifluoroethanol (TFE). Spectra were deconvoluted using the CONTINLL analysis program[3]. Data from reference datasets 4, 7 and SMP180 were averaged to obtain secondary structure estimates[4].

| Peptide | No TFE | | | 30% TFE | | |
|---|---|---|---|---|---|---|
| | α-Helix | β-sheet | Random coil | α-Helix | β-sheet | Random coil |
| | % | | | | | |
| **$UL25^{native(45-94)}$** | 12 | 12 | 76 | 37 | 12 | 51 |
| **$UL25^{scr(45-94)}$** | 8 | 26 | 66 | 31 | 16 | 53 |
| **$UL25^{74-94}$** | 7 | 35 | 58 | 7 | 35 | 58 |
| **$gB^{103-152}$** | 8 | 26 | 66 | 13 | 24 | 63 |
| **$(EAEKAAK)_7EAEK$** | 63 | 3 | 29 | 75 | 2 | 23 |

**Supplementary Table S2. List of primers and gBlock sequence used for cloning procedures described in Materials and Methods.** All primers are listed in the 5'-3' direction. Restriction sites are underlined and mutations are bolded.

| **Primer Name** | **Primer Sequence (5'-3')** | **Restriction Site** |
|---|---|---|
| UL25 rev (JB134) | aaaaaactcgagttattacactgcgctcagatactgagg | XhoI |
| UL25Δ44 Q72A (ED06) | aatgcagcaatg**gcg**gcagcc | Site-directed |
| UL25Δ44 Q72A (ED07) | ggctgc**cgc**cattgctgcatt | Site-directed |
| (EAEKAAK)$_7$EAEK peptide gBlock sequence | aaaaaa ggatcc gaa gcg gaa aaa gcg gcg aaa gaa gcg gaa aaa gcg gcg aaa gaa gcg gaa aaa gaa gcg gaa aaa gcg gcg aaa gaa gcg gaa aaa gcg gcg aaa gaa gcg gaa aaa gaa gcg gaa aaa gcg gcg aaa gaa gcg gaa aaa gcg gcg aaa gaa gcg gaa aaa taa taa ctcgag aaaaaa | BamHI/XhoI |

**Supplementary Table 3. Raw data and background values collected for each reported GUV budding assay.** The reported biological replicate average values (%) are the points presented in Figure 2.

| Biological Replicate | Technical Replicate | Background (BG) | ILVs/GUVs | Raw Value (ILV-BG) | Average (AVG) | ILVs/GUVs | Raw Value (ILV-BG) | Average (AVG) | % Normalized Budding (Raw values/NEC AVG) * 100 | Biological Replicate Average (%) |
|---|---|---|---|---|---|---|---|---|---|---|
| | | | **NEC220** | | | **NEC220 + UL25$^{native(45-94)}$ (1:0.1 NEC:peptide)** | | | | |
| 1 | 1 | 10/100 = 0.10 | 17/100 = 0.17 | 0.07 | 0.09 | 13/100 = 0.13 | 0.03 | 0.10 | 34 | 112 |
| | 2 | 6/100 = 0.06 | 15/100 = 0.15 | 0.09 | | 19/95 = 0.20 | 0.14 | | 161 | |
| | 3 | 8/100 = 0.08 | 18/100 = 0.18 | 0.10 | | 19/95 = 0.20 | 0.12 | | 138 | |
| 2 | 1 | 3/96 = 0.03 | 23/99 = 0.23 | 0.20 | 0.15 | 20/99 = 0.20 | 0.17 | 0.17 | 112 | 111 |
| | 2 | 7/95 = 0.07 | 22/100 = 0.22 | 0.15 | | 26/100 = 0.26 | 0.19 | | 122 | |
| | 3 | 9/100 = 0.09 | 20/100 = 0.20 | 0.11 | | 24/100 =0.24 | 0.15 | | 98 | |
| 3 | 1 | 6/100 = 0.06 | 19/100 = 0.19 | 0.13 | 0.14 | 19/100 = 0.19 | 0.13 | 0.13 | 95 | 95 |
| | 2 | 7/100 = 0.07 | 19/100 = 0.19 | 0.12 | | 19/90 = 0.21 | 0.14 | | 103 | |
| | 3 | 2/100 = 0.02 | 18/100 = 0.18 | 0.16 | | 14/100 = 0.14 | 0.12 | | 88 | |
| | | | **NEC220** | | | **NEC220 + UL25$^{native(45-94)}$ (1:0.3 NEC:peptide)** | | | | |
| 1 | 1 | 3/96 = 0.03 | 23/99 = 0.23 | 0.20 | 0.15 | 15/100 = 0.15 | 0.12 | 0.11 | 78 | 71 |
| | 2 | 7/95 = 0.07 | 22/100 = 0.22 | 0.15 | | 18/100 = 0.18 | 0.11 | | 70 | |
| | 3 | 9/100 = 0.09 | 20/100 = 0.20 | 0.11 | | 19/100 = 0.19 | 0.10 | | 67 | |
| 2 | 1 | 6/100 = 0.06 | 19/100 = 0.19 | 0.13 | 0.14 | 14/100 = 0.14 | 0.08 | 0.9 | 59 | 68 |
| | 2 | 7/100 = 0.07 | 19/100 = 0.19 | 0.12 | | 13/100 = 0.13 | 0.06 | | 44 | |
| | 3 | 2/100 = 0.02 | 18/100 = 0.18 | 0.16 | | 16/100 = 0.16 | 0.14 | | 102 | |
| 3 | 1 | 1/100 = 0.01 | 17/90 = 0.19 | 0.18 | 0.14 | 14/100 = 0.14 | 0.13 | 0.11 | 91 | 75 |
| | 2 | 6/100 = 0.06 | 18/100 = 0.18 | 0.14 | | 14/100 = 0.14 | 0.08 | | 56 | |
| | 3 | 5/100 = 0.05 | 18/100 = 0.18 | 0.13 | | 16/100 = 0.16 | 0.11 | | 77 | |
| | | | **NEC220** | | | **NEC220 + UL25$^{native(45-94)}$ (1:0.6 NEC:peptide)** | | | | |
| 1 | 1 | 8/100 = 0.08 | 19/100 = 0.19 | 0.11 | 0.11 | 16/100 = 0.16 | 0.08 | 0.06 | 71 | 56 |
| | 2 | 9/100 = 0.09 | 18/100 = 0.18 | 0.09 | | 17/100 = 0.17 | 0.08 | | 71 | |
| | 3 | 8/100 = 0.08 | 22/100 = 0.22 | 0.14 | | 11/100 = 0.11 | 0.03 | | 26 | |
| 2 | 1 | 6/100 = 0.06 | 19/100 = 0.19 | 0.13 | 0.14 | 13/95 = 0.14 | 0.08 | 0.08 | 56 | 58 |
| | 2 | 7/100 = 0.07 | 19/100 = 0.19 | 0.12 | | 13/100 = 0.13 | 0.06 | | 44 | |
| | 3 | 2/100 = 0.02 | 18/100 = 0.18 | 0.16 | | 12/100 = 0.12 | 0.10 | | 73 | |
| 3 | 1 | 1/100 = 0.01 | 17/90 = 0.19 | 0.18 | 0.14 | 10/100 = 0.10 | 0.09 | 0.07 | 63 | 49 |
| | 2 | 6/100 = 0.06 | 18/100 = 0.18 | 0.14 | | 11/100 = 0.11 | 0.05 | | 35 | |
| | 3 | 5/100 = 0.05 | 18/100 = 0.18 | 0.13 | | 12/100 = 0.12 | 0.07 | | 49 | |
| | | | **NEC220** | | | **NEC220 + UL25$^{native(45-94)}$ (1:1 NEC:peptide)** | | | | |
| 1 | 1 | 8/100 = 0.08 | 19/100 = 0.19 | 0.11 | 0.11 | 10/100 = 0.10 | 0.02 | 0.04 | 18 | 36 |
| | 2 | 9/100 = 0.09 | 18/100 = 0.18 | 0.09 | | 13/97 = 0.13 | 0.04 | | 39 | |

| Biological Replicate | Technical Replicate | Background (BG) | ILVs/GUVs | Raw Value (ILV-BG) | Average (AVG) | ILVs/GUVs | Raw Value (ILV-BG) | Average (AVG) | % Normalized Budding (Raw values/NEC AVG) * 100 | Biological Replicate Average (%) |
|---|---|---|---|---|---|---|---|---|---|---|
| | 3 | 8/100 = 0.08 | 22/100 = 0.22 | 0.14 | | 14/100 = 0.14 | 0.06 | | 53 | |
| | 1 | 3/90 = 0.03 | 10/81 = 0.12 | 0.09 | | 2/80 = 0.03 | 0.00 | | 9 | |
| 2 | 2 | 3/100 = 0.03 | 12/90 = 0.13 | 0.10 | 0.09 | 2/75 = 0.03 | 0.00 | 0.02 | 3 | 19 |
| | 3 | 3/85 = 0.04 | 9/80 = 0.11 | 0.07 | | 7/80 = 0.09 | 0.07 | | 58 | |
| | 1 | 5/100 = 0.05 | 12/100 = 0.12 | 0.07 | | 7/100 = 0.07 | 0.02 | | 21 | |
| 3 | 2 | 6/100 = 0.06 | 16/105 = 0.15 | 0.09 | 0.10 | 8/100 = 0.08 | 0.02 | 0.03 | 21 | 27 |
| | 3 | 5/100 = 0.05 | 18/100 = 0.18 | 0.13 | | 9/100 = 0.09 | 0.04 | | 41 | |
| | | | **NEC220** | | | **NEC220 + UL25$^{native(45-94)}$ (1:10 NEC:peptide)** | | | | |
| | 1 | 3/96 = 0.03 | 23/99 = 0.23 | 0.20 | | 11/100 = 0.11 | 0.08 | | 52 | |
| 1 | 2 | 7/95 = 0.07 | 22/100 = 0.22 | 0.15 | 0.15 | 7/100 = 0.07 | 0.00 | 0.05 | 0 | 30 |
| | 3 | 9/100 = 0.09 | 20/100 = 0.20 | 0.11 | | 15/100 = 0.15 | 0.06 | | 39 | |
| | 1 | 3/100 = 0.03 | 13/94 = 0.14 | 0.11 | | 3/100 = 0.03 | 0.00 | | 0 | |
| 2 | 2 | 4/100 = 0.04 | 8/90 = 0.09 | 0.05 | 0.07 | 5/91 = 0.05 | 0.01 | 0.02 | 23 | 33 |
| | 3 | 5/100 = 0.05 | 8/91 = 0.09 | 0.04 | | 7/70 = 0.10 | 0.05 | | 77 | |
| | 1 | 5/100 = 0.05 | 12/100 = 0.12 | 0.07 | | 8/100 = 0.08 | 0.03 | | 31 | |
| 3 | 2 | 6/100 = 0.06 | 16/105 = 0.15 | 0.09 | 0.10 | 9/100 = 0.09 | 0.03 | 0.03 | 31 | 34 |
| | 3 | 5/100 = 0.05 | 18/100 = 0.18 | 0.13 | | 9/100 = 0.09 | 0.04 | | 41 | |
| | | | **NEC220** | | | **NEC220 + UL25$^{scr(45-94)}$ (1:0.1 NEC:peptide)** | | | | |
| | 1 | 10/100 = 0.10 | 17/100 = 0.17 | 0.07 | | 21/100 = 0.21 | 0.11 | | 127 | |
| 1 | 2 | 6/100 = 0.06 | 15/100 = 0.15 | 0.09 | 0.09 | 13/100 = 0.13 | 0.07 | 0.10 | 81 | 115 |
| | 3 | 8/100 = 0.08 | 18/100 = 0.18 | 0.10 | | 20/100 = 0.20 | 0.12 | | 138 | |
| | 1 | 3/96 = 0.03 | 23/99 = 0.23 | 0.20 | | 17/100 = 0.17 | 0.14 | | 91 | |
| 2 | 2 | 7/95 = 0.07 | 22/100 = 0.22 | 0.15 | 0.15 | 18/100 = 0.18 | 0.11 | 0.13 | 70 | 84 |
| | 3 | 9/100 = 0.09 | 20/100 = 0.20 | 0.11 | | 23/100 =0.23 | 0.14 | | 92 | |
| | 1 | 6/100 = 0.06 | 19/100 = 0.19 | 0.13 | | 19/96 = 0.20 | 0.14 | | 101 | |
| 3 | 2 | 7/100 = 0.07 | 19/100 = 0.19 | 0.12 | 0.14 | 18/100 = 0.18 | 0.11 | 0.14 | 80 | 104 |
| | 3 | 2/100 = 0.02 | 18/100 = 0.18 | 0.16 | | 20/100 = 0.20 | 0.18 | | 132 | |
| | | | **NEC220** | | | **NEC220 + UL25$^{scr(45-94)}$ (1:0.3 NEC:peptide)** | | | | |
| | 1 | 3/96 = 0.03 | 23/99 = 0.23 | 0.20 | | 20/100 = 0.20 | 0.17 | | 111 | |
| 1 | 2 | 7/95 = 0.07 | 22/100 = 0.22 | 0.15 | 0.15 | 13/100 = 0.13 | 0.08 | 0.10 | 37 | 65 |
| | 3 | 9/100 = 0.09 | 20/100 = 0.20 | 0.11 | | 16/98 = 0.16 | 0.07 | | 48 | |
| | 1 | 6/100 = 0.06 | 19/100 = 0.19 | 0.13 | | 14/90 = 0.16 | 0.10 | | 70 | |
| 2 | 2 | 7/100 = 0.07 | 19/100 = 0.19 | 0.12 | 0.14 | 18/100 = 0.18 | 0.11 | 0.11 | 80 | 82 |
| | 3 | 2/100 = 0.02 | 18/100 = 0.18 | 0.16 | | 15/100 = 0.15 | 0.13 | | 95 | |
| | 1 | 1/100 = 0.01 | 17/90 = 0.19 | 0.18 | | 15/100 = 0.15 | 0.14 | | 98 | |
| 3 | 2 | 6/100 = 0.06 | 18/100 = 0.18 | 0.14 | 0.14 | 15/100 = 0.15 | 0.09 | 0.11 | 63 | 77 |
| | 3 | 5/100 = 0.05 | 18/100 = 0.18 | 0.13 | | 15/100 = 0.15 | 0.10 | | 70 | |

| Biological Replicate | Technical Replicate | Background (BG) | ILVs/GUVs | Raw Value (ILV-BG) | Average (AVG) | ILVs/GUVs | Raw Value (ILV-BG) | Average (AVG) | % Normalized Budding (Raw values/NEC AVG) * 100 | Biological Replicate Average (%) |
|---|---|---|---|---|---|---|---|---|---|---|
| | | | **NEC220** | | | **NEC220 + UL25$^{scr(45-94)}$ (1:0.6 NEC:peptide)** | | | | |
| 1 | 1 | 8/100 = 0.08 | 19/100 = 0.19 | 0.11 | 0.11 | 15/90 = 0.17 | | 0.07 | 76 | 61 |
| | 2 | 9/100 = 0.09 | 18/100 = 0.18 | 0.09 | | 15/100 = 0.15 | 0.06 | | 52 | |
| | 3 | 8/100 = 0.08 | 22/100 = 0.22 | 0.14 | | 14/100 = 0.14 | 0.06 | | 53 | |
| 2 | 1 | 6/100 = 0.06 | 19/100 = 0.19 | 0.13 | 0.14 | 13/100 = 0.13 | 0.07 | 0.08 | 51 | 62 |
| | 2 | 7/100 = 0.07 | 19/100 = 0.19 | 0.12 | | 12/90 = 0.13 | 0.06 | | 46 | |
| | 3 | 2/100 = 0.02 | 18/100 = 0.18 | 0.16 | | 14/100 = 0.14 | 0.12 | | 88 | |
| 3 | 1 | 1/100 = 0.01 | 17/90 = 0.19 | 0.18 | 0.14 | 12/100 = 0.12 | 0.11 | 0.08 | 78 | 56 |
| | 2 | 6/100 = 0.06 | 18/100 = 0.18 | 0.14 | | 11/100 = 0.11 | 0.05 | | 35 | |
| | 3 | 5/100 = 0.05 | 18/100 = 0.18 | 0.13 | | 13/100 = 0.13 | 0.08 | | 56 | |
| | | | **NEC220** | | | **NEC220 + UL25$^{scr(45-94)}$ (1:1 NEC:peptide)** | | | | |
| 1 | 1 | 8/100 = 0.08 | 19/100 = 0.19 | 0.11 | 0.11 | 13/100 = 0.13 | 0.05 | 0.04 | 44 | 38 |
| | 2 | 9/100 = 0.09 | 18/100 = 0.18 | 0.09 | | 13/100 = 0.13 | 0.04 | | 35 | |
| | 3 | 8/100 = 0.08 | 22/100 = 0.22 | 0.14 | | 12/100 = 0.12 | 0.04 | | 35 | |
| 2 | 1 | 3/90 = 0.03 | 10/81 = 0.12 | 0.09 | 0.09 | 7/76 = 0.09 | 0.06 | 0.02 | 65 | 27 |
| | 2 | 3/100 = 0.03 | 12/90 = 0.13 | 0.10 | | 4/100 = 0.04 | 0.01 | | 11 | |
| | 3 | 3/85 = 0.04 | 9/80 = 0.11 | 0.07 | | 4/100 = 0.04 | 0.00 | | 5 | |
| 3 | 1 | 5/100 = 0.05 | 12/100 = 0.12 | 0.07 | 0.10 | 11/100 = 0.11 | 0.06 | 0.02 | 62 | 25 |
| | 2 | 6/100 = 0.06 | 16/105 = 0.15 | 0.09 | | 6/97 = 0.06 | 0.00 | | 2 | |
| | 3 | 5/100 = 0.05 | 18/100 = 0.18 | 0.13 | | 6/100 = 0.06 | 0.01 | | 10 | |
| | | | **NEC220** | | | **NEC220 + UL25$^{scr(45-94)}$ (1:10 NEC:peptide)** | | | | |
| 1 | 1 | 3/96 = 0.03 | 23/99 = 0.23 | 0.20 | 0.15 | 9/95 = 0.09 | 0.06 | 0.05 | 42 | 31 |
| | 2 | 7/95 = 0.07 | 22/100 = 0.22 | 0.15 | | 13/100 = 0.13 | 0.08 | | 37 | |
| | 3 | 9/100 = 0.09 | 20/100 = 0.20 | 0.11 | | 11/100 = 0.11 | 0.02 | | 13 | |
| 2 | 1 | 3/100 = 0.03 | 13/94 = 0.14 | 0.11 | 0.07 | 6/100 = 0.06 | 0.03 | 0.02 | 46 | 29 |
| | 2 | 4/100 = 0.04 | 8/90 = 0.09 | 0.05 | | 5/81 = 0.06 | 0.02 | | 33 | |
| | 3 | 5/100 = 0.05 | 8/91 = 0.09 | 0.04 | | 4/72 = 0.06 | 0.01 | | 9 | |
| 3 | 1 | 5/100 = 0.05 | 12/100 = 0.12 | 0.07 | 0.10 | 9/100 = 0.09 | 0.04 | 0.03 | 41 | 32 |
| | 2 | 6/100 = 0.06 | 16/105 = 0.15 | 0.09 | | 7/100 = 0.07 | 0.01 | | 10 | |
| | 3 | 5/100 = 0.05 | 18/100 = 0.18 | 0.13 | | 9/95 = 0.09 | 0.04 | | 46 | |
| | | | **NEC220** | | | **NEC220 + UL25$^{74-94}$ (1:0.1 NEC:peptide)** | | | | |
| 1 | 1 | 10/100 = 0.10 | 17/100 = 0.17 | 0.07 | 0.09 | 15/100 = 0.15 | 0.05 | 0.08 | 58 | 96 |
| | 2 | 6/100 = 0.06 | 15/100 = 0.15 | 0.09 | | 17/100 = 0.17 | 0.11 | | 127 | |
| | 3 | 8/100 = 0.08 | 18/100 = 0.18 | 0.10 | | 17/100 = 0.17 | 0.09 | | 104 | |
| 2 | 1 | 3/96 = 0.03 | 23/99 = 0.23 | 0.20 | 0.15 | 21/100 = 0.21 | 0.18 | 0.15 | 117 | 97 |

| Biological Replicate | Technical Replicate | Background (BG) | ILVs/GUVs | Raw Value (ILV-BG) | Average (AVG) | ILVs/GUVs | Raw Value (ILV-BG) | Average (AVG) | % Normalized Budding (Raw values/NEC AVG) * 100 | Biological Replicate Average (%) |
|---|---|---|---|---|---|---|---|---|---|---|
| | 2 | 7/95 = 0.07 | 22/100 = 0.22 | 0.15 | | 23/100 = 0.23 | 0.16 | | 102 | |
| | 3 | 9/100 = 0.09 | 20/100 = 0.20 | 0.11 | | 20/100 = 0.20 | 0.11 | | 72 | |
| 3 | 1 | 6/100 = 0.06 | 19/100 = 0.19 | 0.13 | 0.14 | 17/90 = 0.19 | 0.13 | 0.13 | 94 | 97 |
| | 2 | 7/100 = 0.07 | 19/100 = 0.19 | 0.12 | | 21/100 = 0.21 | 0.14 | | 102 | |
| | 3 | 2/100 = 0.02 | 18/100 = 0.18 | 0.16 | | 15/100 = 0.15 | 0.13 | | 95 | |
| | | | **NEC220** | | | **NEC220 + UL25$^{74-94}$ (1:0.3 NEC:peptide)** | | | | |
| 1 | 1 | 3/96 = 0.03 | 23/99 = 0.23 | 0.20 | 0.15 | 25/100 = 0.25 | 0.22 | 0.16 | 143 | 102 |
| | 2 | 7/95 = 0.07 | 22/100 = 0.22 | 0.15 | | 23/100 = 0.23 | 0.16 | | 103 | |
| | 3 | 9/100 = 0.09 | 20/100 = 0.20 | 0.11 | | 18/100 = 0.18 | 0.09 | | 59 | |
| 2 | 1 | 6/100 = 0.06 | 19/100 = 0.19 | 0.13 | 0.14 | 17/100 = 0.17 | 0.11 | 0.13 | 80 | 98 |
| | 2 | 7/100 = 0.07 | 19/100 = 0.19 | 0.12 | | 17/100 = 0.17 | 0.11 | | 73 | |
| | 3 | 2/100 = 0.02 | 18/100 = 0.18 | 0.16 | | 21/100 = 0.21 | 0.19 | | 139 | |
| 3 | 1 | 1/100 = 0.01 | 17/90 = 0.19 | 0.18 | 0.14 | 17/90 = 0.19 | 0.18 | 0.14 | 125 | 98 |
| | 2 | 6/100 = 0.06 | 18/100 = 0.18 | 0.14 | | 17/100 = 0.17 | 0.11 | | 77 | |
| | 3 | 5/100 = 0.05 | 18/100 = 0.18 | 0.13 | | 18/100 = 0.18 | 0.13 | | 91 | |
| | | | **NEC220** | | | **NEC220 + UL25$^{74-94}$ (1:0.6 NEC:peptide)** | | | | |
| 1 | 1 | 6/100 = 0.06 | 19/100 = 0.19 | 0.13 | 0.14 | 18/100 = 0.18 | 0.12 | 0.12 | 88 | 87 |
| | 2 | 7/100 = 0.07 | 19/100 = 0.19 | 0.12 | | 16/95 = 0.17 | 0.10 | | 72 | |
| | 3 | 2/100 = 0.02 | 18/100 = 0.18 | 0.16 | | 16/100 = 0.16 | 0.14 | | 102 | |
| 2 | 1 | 1/100 = 0.01 | 17/90 = 0.19 | 0.18 | 0.14 | 18/93 = 0.19 | 0.18 | 0.15 | 128 | 103 |
| | 2 | 6/100 = 0.06 | 18/100 = 0.18 | 0.14 | | 20/100 = 0.20 | 0.14 | | 98 | |
| | 3 | 5/100 = 0.05 | 18/100 = 0.18 | 0.13 | | 17/100 = 0.17 | 0.12 | | 84 | |
| 3 | 1 | 8/100 = 0.08 | 19/100 = 0.19 | 0.11 | 0.11 | 17/100 = 0.17 | 0.09 | 0.10 | 80 | 90 |
| | 2 | 9/100 = 0.09 | 18/100 = 0.18 | 0.09 | | 20/100 = 0.20 | 0.11 | | 97 | |
| | 3 | 8/100 = 0.08 | 22/100 = 0.22 | 0.14 | | 18/96 = 0.19 | 0.11 | | 95 | |
| | | | **NEC220** | | | **NEC220 + UL25$^{74-94}$ (1:1 NEC:peptide)** | | | | |
| 1 | 1 | 8/100 = 0.08 | 19/100 = 0.19 | 0.11 | 0.11 | 15/96 = 0.16 | 0.08 | 0.09 | 67 | 75 |
| | 2 | 9/100 = 0.09 | 18/100 = 0.18 | 0.09 | | 18/100 = 0.18 | 0.09 | | 79 | |
| | 3 | 8/100 = 0.08 | 22/100 = 0.22 | 0.14 | | 17/100 = 0.17 | 0.09 | | 79 | |
| 2 | 1 | 1/100 = 0.01 | 17/90 = 0.19 | 0.18 | 0.14 | 16/100 = 0.16 | 0.15 | 0.13 | 105 | 93 |
| | 2 | 6/100 = 0.06 | 18/100 = 0.18 | 0.14 | | 18/100 = 0.18 | 0.12 | | 84 | |
| | 3 | 5/100 = 0.05 | 18/100 = 0.18 | 0.13 | | 18/100 = 0.18 | 0.13 | | 91 | |
| 3 | 1 | 5/100 = 0.05 | 12/100 = 0.12 | 0.07 | 0.10 | 10/100 = 0.10 | 0.05 | 0.05 | 51 | 55 |
| | 2 | 6/100 = 0.06 | 16/105 = 0.15 | 0.09 | | 10/100 = 0.10 | 0.04 | | 41 | |
| | 3 | 5/100 = 0.05 | 18/100 = 0.18 | 0.13 | | 12/100 = 0.12 | 0.07 | | 72 | |

| Biological Replicate | Technical Replicate | Background (BG) | ILVs/GUVs | Raw Value (ILV-BG) | Average (AVG) | ILVs/GUVs | Raw Value (ILV-BG) | Average (AVG) | % Normalized Budding (Raw values/NEC AVG) * 100 | Biological Replicate Average (%) |
|---|---|---|---|---|---|---|---|---|---|---|
| | | | **NEC220** | | | **NEC220 + UL25$^{74-94}$ (1:10 NEC:peptide)** | | | | |
| 1 | 1 | 3/96 = 0.03 | 23/99 = 0.23 | 0.20 | 0.15 | 17/100 = 0.17 | 0.14 | 0.10 | 91 | 64 |
| | 2 | 7/95 = 0.07 | 22/100 = 0.22 | 0.15 | | 14/96 = 0.15 | 0.08 | | 47 | |
| | 3 | 9/100 = 0.09 | 20/100 = 0.20 | 0.11 | | 17/100 = 0.17 | 0.08 | | 52 | |
| 2 | 1 | 3/100 = 0.03 | 13/94 = 0.14 | 0.11 | 0.07 | 6/97 = 0.06 | 0.03 | 0.04 | 49 | 66 |
| | 2 | 4/100 = 0.04 | 8/90 = 0.09 | 0.05 | | 6/100 = 0.06 | 0.02 | | 31 | |
| | 3 | 5/100 = 0.05 | 8/91 = 0.09 | 0.04 | | 12/95 = 0.13 | 0.08 | | 117 | |
| 3 | 1 | 5/100 = 0.05 | 12/100 = 0.12 | 0.07 | 0.10 | 15/100 = 0.15 | 0.10 | 0.07 | 103 | 72 |
| | 2 | 6/100 = 0.06 | 16/105 = 0.15 | 0.09 | | 11/100 = 0.11 | 0.05 | | 51 | |
| | 3 | 5/100 = 0.05 | 18/100 = 0.18 | 0.13 | | 11/100 = 0.11 | 0.06 | | 62 | |
| | | | **NEC220** | | | **NEC220 + gB RC$^{103-152}$ (1:0.1 NEC:peptide)** | | | | |
| 1 | 1 | 10/100 = 0.10 | 17/100 = 0.17 | 0.07 | 0.09 | 20/100 = 0.20 | 0.10 | 0.10 | 115 | 115 |
| | 2 | 6/100 = 0.06 | 15/100 = 0.15 | 0.09 | | 17/100 = 0.17 | 0.11 | | 127 | |
| | 3 | 8/100 = 0.08 | 18/100 = 0.18 | 0.10 | | 17/100 = 0.17 | 0.09 | | 104 | |
| 2 | 1 | 3/96 = 0.03 | 23/99 = 0.23 | 0.20 | 0.15 | 22/100 = 0.22 | 0.19 | 0.18 | 124 | 120 |
| | 2 | 7/95 = 0.07 | 22/100 = 0.22 | 0.15 | | 25/98 = 0.26 | 0.19 | | 119 | |
| | 3 | 9/100 = 0.09 | 20/100 = 0.20 | 0.11 | | 24/90 = 0.27 | 0.18 | | 116 | |
| 3 | 1 | 6/100 = 0.06 | 19/100 = 0.19 | 0.13 | 0.14 | 19/90 = 0.21 | 0.15 | 0.15 | 111 | 107 |
| | 2 | 7/100 = 0.07 | 19/100 = 0.19 | 0.12 | | 16/100 = 0.16 | 0.09 | | 66 | |
| | 3 | 2/100 = 0.02 | 18/100 = 0.18 | 0.16 | | 21/97 = 0.22 | 0.20 | | 144 | |
| | | | **NEC220** | | | **NEC220 + gB RC$^{103-152}$ (1:0.3 NEC:peptide)** | | | | |
| 1 | 1 | 3/96 = 0.03 | 23/99 = 0.23 | 0.20 | 0.15 | 21/100 = 0.21 | 0.18 | 0.16 | 117 | 102 |
| | 2 | 7/95 = 0.07 | 22/100 = 0.22 | 0.15 | | 23/100 = 0.23 | 0.16 | | 103 | |
| | 3 | 9/100 = 0.09 | 20/100 = 0.20 | 0.11 | | 22/100 = 0.22 | 0.13 | | 85 | |
| 2 | 1 | 6/100 = 0.06 | 19/100 = 0.19 | 0.13 | 0.14 | 18/100 = 0.18 | 0.12 | 0.14 | 88 | 105 |
| | 2 | 7/100 = 0.07 | 19/100 = 0.19 | 0.12 | | 21/100 = 0.21 | 0.14 | | 102 | |
| | 3 | 2/100 = 0.02 | 18/100 = 0.18 | 0.16 | | 19/100 = 0.19 | 0.17 | | 124 | |
| 3 | 1 | 1/100 = 0.01 | 17/90 = 0.19 | 0.18 | 0.14 | 17/100 = 0.17 | 0.16 | 0.14 | 112 | 96 |
| | 2 | 6/100 = 0.06 | 18/100 = 0.18 | 0.14 | | 18/100 = 0.18 | 0.12 | | 84 | |
| | 3 | 5/100 = 0.05 | 18/100 = 0.18 | 0.13 | | 17/100 = 0.17 | 0.12 | | 91 | |
| | | | **NEC220** | | | **NEC220 + gB RC$^{103-152}$ (1:0.6 NEC:peptide)** | | | | |
| 1 | 1 | 6/100 = 0.06 | 19/100 = 0.19 | 0.13 | 0.14 | 16/100 = 0.16 | 0.10 | 0.12 | 73 | 85 |
| | 2 | 7/100 = 0.07 | 19/100 = 0.19 | 0.12 | | 14/100 = 0.14 | 0.07 | | 51 | |
| | 3 | 2/100 = 0.02 | 18/100 = 0.18 | 0.16 | | 20/100 = 0.20 | 0.18 | | 132 | |
| 2 | 1 | 1/100 = 0.01 | 17/90 = 0.19 | 0.18 | 0.14 | 16/100 = 0.16 | 0.15 | 0.13 | 105 | 93 |
| | 2 | 6/100 = 0.06 | 18/100 = 0.18 | 0.14 | | 19/100 = 0.19 | 0.13 | | 91 | |

| Biological Replicate | Technical Replicate | Background (BG) | ILVs/GUVs | Raw Value (ILV-BG) | Average (AVG) | ILVs/GUVs | Raw Value (ILV-BG) | Average (AVG) | % Normalized Budding (Raw values/NEC AVG) * 100 | Biological Replicate Average (%) |
|---|---|---|---|---|---|---|---|---|---|---|
| | 3 | 5/100 = 0.05 | 18/100 = 0.18 | 0.13 | | 16/94 = 0.17 | 0.12 | | 84 | |
| 3 | 1 | 8/100 = 0.08 | 19/100 = 0.19 | 0.11 | 0.11 | 16/100 = 0.16 | 0.08 | 0.10 | 71 | 91 |
| | 2 | 9/100 = 0.09 | 18/100 = 0.18 | 0.09 | | 18/100 = 0.18 | 0.09 | | 80 | |
| | 3 | 8/100 = 0.08 | 22/100 = 0.22 | 0.14 | | 22/100 = 0.22 | 0.14 | | 124 | |
| | | | **NEC220** | | | **NEC220 + gB RC$^{103-152}$ (1:1 NEC:peptide)** | | | | |
| 1 | 1 | 8/100 = 0.08 | 19/100 = 0.19 | 0.11 | 0.11 | 18/95 = 0.19 | 0.11 | 0.12 | 97 | 103 |
| | 2 | 9/100 = 0.09 | 18/100 = 0.18 | 0.09 | | 19/100 = 0.19 | 0.10 | | 88 | |
| | 3 | 8/100 = 0.08 | 22/100 = 0.22 | 0.14 | | 22/100 = 0.22 | 0.14 | | 124 | |
| 2 | 1 | 1/100 = 0.01 | 17/90 = 0.19 | 0.18 | 0.14 | 17/100 = 0.17 | 0.16 | 0.14 | 112 | 98 |
| | 2 | 6/100 = 0.06 | 18/100 = 0.18 | 0.14 | | 16/100 = 0.16 | 0.10 | | 70 | |
| | 3 | 5/100 = 0.05 | 18/100 = 0.18 | 0.13 | | 19/90 = 0.21 | 0.16 | | 113 | |
| 3 | 1 | 5/100 = 0.05 | 12/100 = 0.12 | 0.07 | 0.10 | 13/100 = 0.13 | 0.08 | 0.08 | 82 | 82 |
| | 2 | 6/100 = 0.06 | 16/105 = 0.15 | 0.09 | | 15/100 = 0.15 | 0.09 | | 92 | |
| | 3 | 5/100 = 0.05 | 18/100 = 0.18 | 0.13 | | 12/100 = 0.12 | 0.07 | | 72 | |
| | | | **NEC220** | | | **NEC220 + gB RC$^{103-152}$ (1:10 NEC:peptide)** | | | | |
| 1 | 1 | 3/96 = 0.03 | 23/99 = 0.23 | 0.20 | 0.15 | 20/91 = 0.22 | 0.19 | 0.12 | 124 | 82 |
| | 2 | 7/95 = 0.07 | 22/100 = 0.22 | 0.15 | | 19/100 = 0.19 | 0.12 | | 76 | |
| | 3 | 9/100 = 0.09 | 20/100 = 0.20 | 0.11 | | 16/100 = 0.16 | 0.07 | | 46 | |
| 2 | 1 | 3/100 = 0.03 | 13/94 = 0.14 | 0.11 | 0.07 | 9/90 = 0.10 | 0.07 | 0.05 | 108 | 73 |
| | 2 | 4/100 = 0.04 | 8/90 = 0.09 | 0.05 | | 8/90 = 0.09 | 0.05 | | 75 | |
| | 3 | 5/100 = 0.05 | 8/91 = 0.09 | 0.04 | | 5/67 = 0.07 | 0.02 | | 38 | |
| 3 | 1 | 5/100 = 0.05 | 12/100 = 0.12 | 0.07 | 0.10 | 9/97 = 0.09 | 0.04 | 0.06 | 44 | 63 |
| | 2 | 6/100 = 0.06 | 16/105 = 0.15 | 0.09 | | 12/100 = 0.12 | 0.06 | | 62 | |
| | 3 | 5/100 = 0.05 | 18/100 = 0.18 | 0.13 | | 13/100 = 0.13 | 0.08 | | 82 | |
| | | | **NEC220** | | | **NEC220 + $(EAEKAAK)_7$EAEK (1:0.1 NEC:peptide)** | | | | |
| 1 | 1 | 3/96 = 0.03 | 23/99 = 0.23 | 0.20 | 0.15 | 22/100 = 0.22 | 0.19 | 0.15 | 124 | 97 |
| | 2 | 7/95 = 0.07 | 22/100 = 0.22 | 0.15 | | 24/100 = 0.24 | 0.17 | | 109 | |
| | 3 | 9/100 = 0.09 | 20/100 = 0.20 | 0.11 | | 18/100 = 0.18 | 0.09 | | 59 | |
| 2 | 1 | 6/100 = 0.06 | 19/100 = 0.19 | 0.13 | 0.14 | 21/93 = 0.23 | 0.17 | 0.15 | 121 | 109 |
| | 2 | 7/100 = 0.07 | 19/100 = 0.19 | 0.12 | | 19/100 = 0.19 | 0.12 | | 88 | |
| | 3 | 2/100 = 0.02 | 18/100 = 0.18 | 0.16 | | 18/100 = 0.18 | 0.16 | | 117 | |
| 3 | 1 | 8/100 = 0.08 | 19/100 = 0.19 | 0.11 | 0.11 | 17/100 = 0.17 | 0.09 | 0.09 | 80 | 82 |
| | 2 | 9/100 = 0.09 | 18/100 = 0.18 | 0.09 | | 16/100 = 0.16 | 0.07 | | 62 | |
| | 3 | 8/100 = 0.08 | 22/100 = 0.22 | 0.14 | | 20/100 = 0.20 | 0.12 | | 106 | |
| | | | **NEC220** | | | **NEC220 + $(EAEKAAK)_7$EAEK (1:0.3 NEC:peptide)** | | | | |
| 1 | 1 | 3/96 = 0.03 | 23/99 = 0.23 | 0.20 | 0.15 | 23/100 = 0.23 | 0.20 | 0.15 | 130 | 99 |

| Biological Replicate | Technical Replicate | Background (BG) | ILVs/GUVs | Raw Value (ILV-BG) | Average (AVG) | ILVs/GUVs | Raw Value (ILV-BG) | Average (AVG) | % Normalized Budding (Raw values/NEC AVG) * 100 | Biological Replicate Average (%) |
|---|---|---|---|---|---|---|---|---|---|---|
| | 2 | 7/95 = 0.07 | 22/100 = 0.22 | 0.15 | | 20/100 = 0.20 | 0.13 | | 83 | |
| | 3 | 9/100 = 0.09 | 20/100 = 0.20 | 0.11 | | 21/96 = 0.22 | 0.13 | | 84 | |
| | 1 | 6/100 = 0.06 | 19/100 = 0.19 | 0.13 | | 20/100 = 0.20 | 0.14 | | 102 | |
| 2 | 2 | 7/100 = 0.07 | 19/100 = 0.19 | 0.12 | 0.14 | 18/100 = 0.18 | 0.11 | 0.13 | 80 | 95 |
| | 3 | 2/100 = 0.02 | 18/100 = 0.18 | 0.16 | | 16/100 = 0.16 | 0.14 | | 102 | |
| | 1 | 1/100 = 0.01 | 17/90 = 0.19 | 0.18 | | 17/100 = 0.17 | 0.16 | | 112 | |
| 3 | 2 | 6/100 = 0.06 | 18/100 = 0.18 | 0.14 | 0.14 | 16/100 = 0.16 | 0.10 | 0.14 | 70 | 96 |
| | 3 | 5/100 = 0.05 | 18/100 = 0.18 | 0.13 | | 20/100 = 0.20 | 0.15 | | 105 | |
| | | | **NEC220** | | | **NEC220 + (EAEKAAK)$_7$EAEK (1:0.6 NEC:peptide)** | | | | |
| | 1 | 6/100 = 0.06 | 19/100 = 0.19 | 0.13 | | 18/100 = 0.18 | 0.12 | | 88 | |
| 1 | 2 | 7/100 = 0.07 | 19/100 = 0.19 | 0.12 | 0.14 | 15/100 = 0.15 | 0.08 | 0.12 | 58 | 88 |
| | 3 | 2/100 = 0.02 | 18/100 = 0.18 | 0.16 | | 18/100 = 0.18 | 0.16 | | 117 | |
| | 1 | 1/100 = 0.01 | 17/90 = 0.19 | 0.18 | | 19/100 = 0.19 | 0.18 | | 126 | |
| 2 | 2 | 6/100 = 0.06 | 18/100 = 0.18 | 0.14 | 0.14 | 21/100 = 0.21 | 0.15 | 0.16 | 105 | 112 |
| | 3 | 5/100 = 0.05 | 18/100 = 0.18 | 0.13 | | 20/100 = 0.20 | 0.15 | | 105 | |
| | 1 | 8/100 = 0.08 | 19/100 = 0.19 | 0.11 | | 16/95 = 0.17 | 0.09 | | 78 | |
| 3 | 2 | 9/100 = 0.09 | 18/100 = 0.18 | 0.09 | 0.11 | 15/100 = 0.15 | 0.06 | 0.10 | 53 | 85 |
| | 3 | 8/100 = 0.08 | 22/100 = 0.22 | 0.14 | | 22/100 = 0.22 | 0.14 | | 124 | |
| | | | **NEC220** | | | **NEC220 + (EAEKAAK)$_7$EAEK (1:1 NEC:peptide)** | | | | |
| | 1 | 8/100 = 0.08 | 19/100 = 0.19 | 0.11 | | 19/90 = 0.21 | 0.13 | | 116 | |
| 1 | 2 | 9/100 = 0.09 | 18/100 = 0.18 | 0.09 | 0.11 | 21/100 = 0.21 | 0.12 | 0.12 | 106 | 106 |
| | 3 | 8/100 = 0.08 | 22/100 = 0.22 | 0.14 | | 19/100 = 0.19 | 0.08 | | 97 | |
| | 1 | 6/100 = 0.06 | 19/100 = 0.19 | 0.13 | | 18/100 = 0.18 | 0.12 | | 88 | |
| 2 | 2 | 7/100 = 0.07 | 19/100 = 0.19 | 0.12 | 0.14 | 17/100 = 0.17 | 0.10 | 0.12 | 73 | 90 |
| | 3 | 2/100 = 0.02 | 18/100 = 0.18 | 0.16 | | 17/100 = 0.17 | 0.15 | | 110 | |
| | 1 | 1/100 = 0.01 | 17/90 = 0.19 | 0.18 | | 15/90 = 0.17 | 0.16 | | 110 | |
| 3 | 2 | 6/100 = 0.06 | 18/100 = 0.18 | 0.14 | 0.14 | 18/100 = 0.18 | 0.12 | 0.16 | 84 | 102 |
| | 3 | 5/100 = 0.05 | 18/100 = 0.18 | 0.13 | | 21/100 = 0.21 | 0.16 | | 112 | |
| | | | **NEC220** | | | **NEC220 + (EAEKAAK)$_7$EAEK (1:10 NEC:peptide)** | | | | |
| | 1 | 9/100 = 0.09 | 0.04 | 0.05 | | 19/99 = 0.19 | 0.15 | | 34 | |
| 1 | 2 | 11/100 = 0.11 | 0.04 | 0.07 | 0.06 | 16/100 = 0.16 | 0.12 | 0.15 | 45 | 40 |
| | 3 | 10/100 = 0.10 | 0.04 | 0.06 | | 21/100 = 0.21 | 0.17 | | 41 | |
| | 1 | 15/101 = 0.15 | 0.06 | 0.09 | | 26/100 = 0.26 | 0.20 | | 41 | |
| 2 | 2 | 19/98 = 0.19 | 0.06 | 0.13 | 0.11 | 22/100 = 0.22 | 0.16 | 0.21 | 63 | 53 |
| | 3 | 14/100 = 0.14 | 0.02 | 0.12 | | 30/100 = 0.30 | 0.28 | | 56 | |
| 3 | 1 | 9/100 = 0.09 | 0.04 | 0.05 | 0.08 | 17/97 = 0.18 | 0.14 | 0.14 | 35 | 54 |
| | 2 | 12/100 = 0.12 | 0.04 | 0.08 | | 16/96 = 0.16 | 0.12 | | 57 | |

| Biological Replicate | Technical Replicate | Background (BG) | ILVs/GUVs | Raw Value (ILV-BG) | Average (AVG) | ILVs/GUVs | Raw Value (ILV-BG) | Average (AVG) | % Normalized Budding (Raw values/NEC AVG) * 100 | Biological Replicate Average (%) |
|---|---|---|---|---|---|---|---|---|---|---|
| | 3 | 13/100 = 0.13 | 0.03 | 0.10 | | 19/100 = 0.19 | 0.13 | | 71 | |
| | | | **NEC220** | | | **NEC220 + UL25Δ44 Q72A (1:0.1 NEC:UL25)** | | | | |
| 1 | 1 | 10/100 = 0.10 | 17/100 = 0.17 | 0.07 | 0.09 | 14/100 = 0.14 | 0.04 | 0.09 | 46 | 107 |
| | 2 | 6/100 = 0.06 | 15/100 = 0.15 | 0.09 | | 17/90 = 0.19 | 0.13 | | 149 | |
| | 3 | 8/100 = 0.08 | 18/100 = 0.18 | 0.10 | | 19/100 = 0.19 | 0.11 | | 127 | |
| 2 | 1 | 3/96 = 0.03 | 23/99 = 0.23 | 0.20 | 0.15 | 21/100 = 0.21 | 0.18 | 0.17 | 117 | 110 |
| | 2 | 7/95 = 0.07 | 22/100 = 0.22 | 0.15 | | 28/100 = 0.28 | 0.21 | | 135 | |
| | 3 | 9/100 = 0.09 | 20/100 = 0.20 | 0.11 | | 21/100 = 0.21 | 0.12 | | 79 | |
| 3 | 1 | 6/100 = 0.06 | 19/100 = 0.19 | 0.13 | 0.14 | 17/100 = 0.17 | 0.11 | 0.13 | 80 | 97 |
| | 2 | 7/100 = 0.07 | 19/100 = 0.19 | 0.12 | | 14/96 = 0.15 | 0.08 | | 55 | |
| | 3 | 2/100 = 0.02 | 18/100 = 0.18 | 0.16 | | 23/100 = 0.23 | 0.21 | | 154 | |
| | | | **NEC220** | | | **NEC220 + UL25Δ44 Q72A (1:0.3 NEC:UL25)** | | | | |
| 1 | 1 | 3/96 = 0.03 | 23/99 = 0.23 | 0.20 | 0.15 | 27/97 = 0.28 | 0.25 | 0.15 | 162 | 101 |
| | 2 | 7/95 = 0.07 | 22/100 = 0.22 | 0.15 | | 17/100 = 0.17 | 0.10 | | 63 | |
| | 3 | 9/100 = 0.09 | 20/100 = 0.20 | 0.11 | | 21/100 = 0.21 | 0.11 | | 79 | |
| 2 | 1 | 6/100 = 0.06 | 19/100 = 0.19 | 0.13 | 0.14 | 16/100 = 0.16 | 0.10 | 0.12 | 73 | 88 |
| | 2 | 7/100 = 0.07 | 19/100 = 0.19 | 0.12 | | 16/100 = 0.16 | 0.09 | | 66 | |
| | 3 | 2/100 = 0.02 | 18/100 = 0.18 | 0.16 | | 19/100 = 0.19 | 0.17 | | 124 | |
| 3 | 1 | 1/100 = 0.01 | 17/90 = 0.19 | 0.18 | 0.14 | 21/100 = 0.21 | 0.20 | 0.14 | 140 | 98 |
| | 2 | 6/100 = 0.06 | 18/100 = 0.18 | 0.14 | | 15/100 = 0.15 | 0.09 | | 63 | |
| | 3 | 5/100 = 0.05 | 18/100 = 0.18 | 0.13 | | 18/100 = 0.18 | 0.13 | | 91 | |
| | | | **NEC220** | | | **NEC220 + UL25Δ44 Q72A (1:0.6 NEC:UL25)** | | | | |
| 1 | 1 | 8/100 = 0.08 | 19/100 = 0.19 | 0.11 | 0.11 | 16/100 = 0.16 | 0.08 | 0.11 | 71 | 97 |
| | 2 | 9/100 = 0.09 | 18/100 = 0.18 | 0.09 | | 21/100 = 0.21 | 0.12 | | 106 | |
| | 3 | 8/100 = 0.08 | 22/100 = 0.22 | 0.14 | | 21/100 = 0.21 | 0.13 | | 115 | |
| 2 | 1 | 6/100 = 0.06 | 19/100 = 0.19 | 0.13 | 0.14 | 18/100 = 0.18 | 0.08 | 0.13 | 88 | 93 |
| | 2 | 7/100 = 0.07 | 19/100 = 0.19 | 0.12 | | 16/94 = 0.17 | 0.10 | | 73 | |
| | 3 | 2/100 = 0.02 | 18/100 = 0.18 | 0.16 | | 18/100 = 0.18 | 0.16 | | 117 | |
| 3 | 1 | 1/100 = 0.01 | 17/90 = 0.19 | 0.18 | 0.14 | 20/100 = 0.20 | 0.19 | 0.15 | 133 | 105 |
| | 2 | 6/100 = 0.06 | 18/100 = 0.18 | 0.14 | | 15/90 = 0.17 | 0.11 | | 75 | |
| | 3 | 5/100 = 0.05 | 18/100 = 0.18 | 0.13 | | 19/93 = 0.20 | 0.15 | | 108 | |
| | | | **NEC220** | | | **NEC220 + UL25Δ44 Q72A (1:1 NEC:UL25)** | | | | |
| 1 | 1 | 8/100 = 0.08 | 19/100 = 0.19 | 0.11 | 0.11 | 19/91 = 0.21 | 0.13 | 0.13 | 113 | 112 |
| | 2 | 9/100 = 0.09 | 18/100 = 0.18 | 0.09 | | 21/80 = 0.27 | 0.18 | | 152 | |
| | 3 | 8/100 = 0.08 | 22/100 = 0.22 | 0.14 | | 16/100 = 0.16 | 0.08 | | 71 | |
| 2 | 1 | 1/100 = 0.01 | 17/90 = 0.19 | 0.18 | 0.14 | 16/100 = 0.16 | 0.15 | 0.15 | 105 | 105 |

| Biological Replicate | Technical Replicate | Background (BG) | ILVs/GUVs | Raw Value (ILV-BG) | Average (AVG) | ILVs/GUVs | Raw Value (ILV-BG) | Average (AVG) | % Normalized Budding (Raw values/NEC AVG) * 100 | Biological Replicate Average (%) |
|---|---|---|---|---|---|---|---|---|---|---|
| | 2 | 6/100 = 0.06 | 18/100 = 0.18 | 0.14 | | 22/100 = 0.22 | 0.16 | | 112 | |
| | 3 | 5/100 = 0.05 | 18/100 = 0.18 | 0.13 | | 19/100 = 0.19 | 0.14 | | 98 | |
| 3 | 1 | 3/90 = 0.03 | 10/81 = 0.12 | 0.09 | 0.09 | 11/97 = 0.11 | 0.08 | 0.09 | 89 | 100 |
| | 2 | 3/100 = 0.03 | 12/90 = 0.13 | 0.10 | | 11/90 = 0.12 | 0.09 | | 102 | |
| | 3 | 3/85 = 0.04 | 9/80 = 0.11 | 0.07 | | 13/97 = 0.13 | 0.09 | | 109 | |
| | | | **NEC220** | | | **NEC220 + UL25Δ44 Q72A (1:10 NEC:UL25)** | | | | |
| 1 | 1 | 3/100 = 0.03 | 13/94 = 0.14 | 0.11 | 0.07 | 4/100 = 0.04 | 0.01 | 0.00 | 15 | 12 |
| | 2 | 4/100 = 0.04 | 8/90 = 0.09 | 0.05 | | 4/100 = 0.04 | 0.00 | | 0 | |
| | 3 | 5/100 = 0.05 | 8/91 = 0.09 | 0.04 | | 5/80 = 0.06 | 0.01 | | 19 | |
| 2 | 1 | 5/100 = 0.05 | 12/100 = 0.12 | 0.07 | 0.10 | 5/100 = 0.05 | 0.00 | 0.00 | 0 | 7 |
| | 2 | 6/100 = 0.06 | 16/105 = 0.15 | 0.09 | | 6/97 = 0.06 | 0.00 | | 2 | |
| | 3 | 5/100 = 0.05 | 18/100 = 0.18 | 0.13 | | 7/100 = 0.07 | 0.02 | | 21 | |
| 3 | 1 | 3/87 = 0.03 | 17/85 = 0.20 | 0.17 | 0.14 | 3/85 = 0.03 | 0.00 | 0.01 | 0 | 7 |
| | 2 | 2/96 = 0.02 | 15/94 = 0.16 | 0.14 | | 5/101 = 0.05 | 0.03 | | 20 | |
| | 3 | 3/103 = 0.03 | 15/97= 0.15 | 0.12 | | 3/95 = 0.03 | 0.00 | | 2 | |

### References


1. Gautier, R., Douguet, D., Antonny, B. & Drin, G. HELIQUEST: a web server to screen sequences with specific alpha-helical properties. *Bioinformatics (Oxford, England)*. **24**, 2101-2102, doi:10.1093/bioinformatics/btn392 (2008).

2. Whitmore, L. & Wallace, B. A. Protein secondary structure analyses from circular dichroism spectroscopy: methods and reference databases. *Biopolymers*. **89**, 392-400, doi:10.1002/bip.20853 (2008).

3. van Stokkum, I. H., Spoelder, H. J., Bloemendal, M., van Grondelle, R. & Groen, F. C. Estimation of protein secondary structure and error analysis from circular dichroism spectra. *Anal. Biochem*. **191**, 110-118 (1990).

4. Sreerama, N. & Woody, R. W. Estimation of protein secondary structure from circular dichroism spectra: comparison of CONTIN, SELCON, and CDSSTR methods with an expanded reference set. *Anal. Biochem*. **287**, 252-260, doi:10.1006/abio.2000.4880 (2000).
